# Distinct cellular expression and subcellular localization of Kv2 voltage-gated K^+^ channel subtypes in dorsal root ganglion neurons conserved between mice and humans

**DOI:** 10.1101/2023.03.01.530679

**Authors:** Robert G. Stewart, Miriam Camacena, Bryan A. Copits, Jon T. Sack

**Affiliations:** Department of Physiology and Membrane Biology, University of California Davis, Davis, CA 95616, USA; Washington University Pain Center, Washington University School of Medicine, St. Louis, MO 63110, USA; Department of Anesthesiology, Washington University School of Medicine, St. Louis, MO 63110, USA; Department of Anesthesiology and Pain Medicine, University of California Davis, Davis, CA 95616, USA

**Keywords:** Dorsal root ganglion, Voltage gated ion channels, Kv2.1, Kv2.2, Somatosensory neurons

## Abstract

The distinct organization of Kv2 voltage-gated potassium channels on and near the cell body of brain neurons enables their regulation of action potentials and specialized membrane contact sites. Somatosensory neurons have a pseudounipolar morphology and transmit action potentials from peripheral nerve endings through axons that bifurcate to the spinal cord and the cell body within ganglia including the dorsal root ganglia (DRG). Kv2 channels regulate action potentials in somatosensory neurons, yet little is known about where Kv2 channels are located. Here we define the cellular and subcellular localization of the Kv2 paralogs, Kv2.1 and Kv2.2, in DRG somatosensory neurons with a panel of antibodies, cell markers, and genetically modified mice. We find that relative to spinal cord neurons, DRG neurons have similar levels of detectable Kv2.1, and higher levels of Kv2.2. In older mice, detectable Kv2.2 remains similar while detectable Kv2.1 decreases. Both Kv2 subtypes adopt clustered subcellular patterns that are distinct from central neurons. Most DRG neurons co-express Kv2.1 and Kv2.2, although neuron subpopulations show preferential expression of Kv2.1 or Kv2.2. We find that Kv2 protein expression and subcellular localization is similar between mouse and human DRG neurons. We conclude that the organization of both Kv2 channels is consistent with physiological roles in the somata and stem axons of DRG neurons. The general prevalence of Kv2.2 in DRG as compared to central neurons and the enrichment of Kv2.2 relative to detectable Kv2.1, in older mice, proprioceptors, and axons suggest more widespread roles for Kv2.2 in DRG neurons.

**Significance statement:** The subcellular distribution of Kv2 voltage-gated potassium channels enable compartment-specific modulation of membrane excitability and organization of membrane contact sites. Here we identify subcellular distributions of the Kv2 paralogs, Kv2.1 and Kv2.2, in somatosensory neurons that bear similarities to and distinctions from central neurons. The distribution of Kv2 channels is similar in mouse and human somatosensory neurons. These results identify unique locations of Kv2 channels in somatosensory neurons that could enable roles in sensory information processing.

## INTRODUCTION

The subcellular localization of voltage-gated ion channels determines how electrical signals are propagated. The two members of the Kv2 family of voltage gated potassium channels, Kv2.1 and Kv2.2, are important for modulating electrical signals in mammalian somatosensory neurons (Bocksteins et al., 2009; Lee et al., 2020; Sun et al., 2022; Tsantoulas et al., 2014; Zheng et al., 2019), yet little is known about where Kv2 channels are localized in these neurons. In central neurons, Kv2 channels are sequestered to specific subcellular regions and identification of where these channels are has helped elucidate their functional roles (Bishop et al., 2015; Du, Tao-Cheng, Zerfas, & McBain, 1998; Irie, 2021; Jensen et al., 2017; Johnson et al., 2018; Kihira, Hermanstyne, & Misonou, 2010; Kirmiz, Vierra, Palacio, & Trimmer, 2018; Misonou, Mohapatra, Menegola, & Trimmer, 2005; Muennich & Fyffe, 2004; Romer et al., 2014; Scannevin, Murakoshi, Rhodes, & Trimmer, 1996; Trimmer, 1991; Vierra, O’Dwyer, Matsumoto, Santana, & Trimmer, 2021). Establishing where Kv2 channels are in somatosensory neurons likewise identifies sites where they could function.

Among vertebrates, the Kv2 family contains two conserved paralogs, Kv2.1 and Kv2.2, which can assemble into homo- or hetero-tetramers to form voltage gated K^+^ channels (Blaine & Ribera, 1998; Kihira et al., 2010). In central neurons, Kv2.1 and Kv2.2 channels have unique cellular expression, subcellular localization, and show distinct physiological roles (Bishop et al., 2015; Newkirk et al., 2022). Among cortical neuron types, Kv2.1 is more broadly distributed than Kv2.2 (Bishop et al., 2015). Little is known of how Kv2.1 or Kv2.2 channels are distributed among somatosensory neuron subtypes. In central neurons, Kv2.1 and Kv2.2 channels localize in clusters on the cell soma, proximal dendrites, and axon initial segment (Bishop et al., 2015; King, Manning, & Trimmer, 2014; Sarmiere, Weigle, & Tamkun, 2008; Scannevin et al., 1996; Trimmer, 1991). Clustered Kv2 channels form endoplasmic reticulum-plasma membrane junctions (Johnson et al., 2018; Kirmiz et al., 2018), where they perform nonconducting functions which include mediating coupling of L-type calcium channels to ryanodine receptors (Vierra et al., 2021), and Ca^2+^ uptake (Panzera et al., 2022). In somatosensory neurons it remains unknown whether Kv2 channels form organized structures or are localized to specific subcellular compartments. Identifying if channel expression is specific to neuron subtypes and within specific subcellular domains could indicate whether Kv2.1 and Kv2.2 channels also have distinct physiological roles in somatosensory neurons.

Kv2 channels are expressed and play important functional roles in DRG. Kv2 transcripts have been detected in all classes of DRG neurons with a notable absence of Kv2.1 mRNA in proprioceptors (Usoskin et al., 2015; Wangzhou et al., 2020; Zheng et al., 2019). Presence of Kv2.1 protein in DRG of mice and rats has been identified by western blot and immunohistochemistry respectively (Sun et al., 2022; Tsantoulas et al., 2012). Pharmacological inhibition of Kv2 channels enhances rat DRG neuron responsiveness to sustained inputs (Tsantoulas et al., 2014). Kv2 conductances have been identified in several mouse DRG neuron subtypes (Bocksteins et al., 2009; Regnier, Bocksteins, Van de Vijver, Snyders, & van Bogaert, 2016; Zheng et al., 2019). Kv2 channels have also been implicated in nociception. Kv2.1 and Kv2.2 mRNA transcript levels are decreased after peripheral axotomy of rat DRG neurons (Tsantoulas et al., 2014). Suppression of Kv2.1 transcription by the epigenetic factor Cdyl is implicated in regulating pain sensation (Sun et al., 2022). A missing piece of the puzzle of how Kv2 channels are involved in somatosensory neuron function is the location of the channels themselves. Here we use antibodies against Kv2.1 and Kv2.2 to define their cellular expression and subcellular localization in DRG neurons of both mice and humans.

## MATERIALS AND METHODS

### Mice

This study was approved by the UC Davis Institutional Animal Care and Use Committee and conforms to guidelines established by the NIH. Mice were maintained on a 12 h light/dark cycle, and food and water was provided *ad libitum.* Kv2.1 KO and Kv2.2 KO mice were from breeding Kv2.1+/- or Kv2.2+/- heterozygous mice such that WT mice were from breeding pairs that generated KO mice. Kv2.1 and Kv2.2 DKO mice were generated from breeding Kv2.1+/- heterozygous and Kv2.2-/- homozygous mice. WT controls for Kv2.1/Kv2.2 DKO experiments were C57BL/6J mice purchased from Jackson Laboratory (stock #000664). The *PV^Ai14^* mouse line was a generous gift from Dr. Theanne Griffith (University of California Davis, Davis CA) and were a cross of *Rosa26^Ai14^* (stock #007914, MGI: J:155793) and *PVcre* (stock #008069, MGI:J: 100886) mice. The *MrgprD^GFP^* mouse line was a generous gift from Dr. David Ginty (Harvard University, Boston MA) (MGI: 3521853). Detailed information about the genotype, age, sex and level of spinal column of mice used in each figure can be found in Table 1.

**Table 1.**
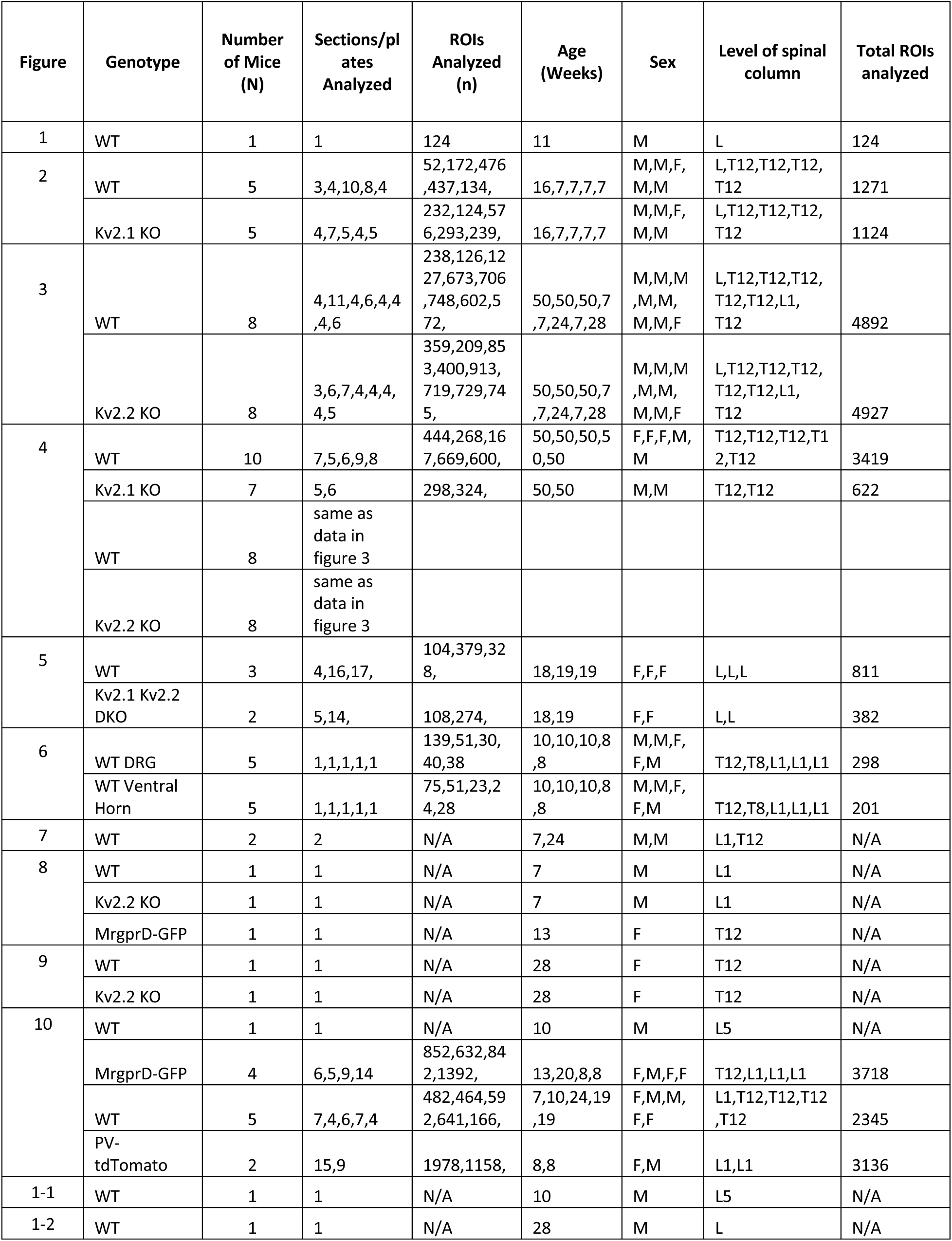
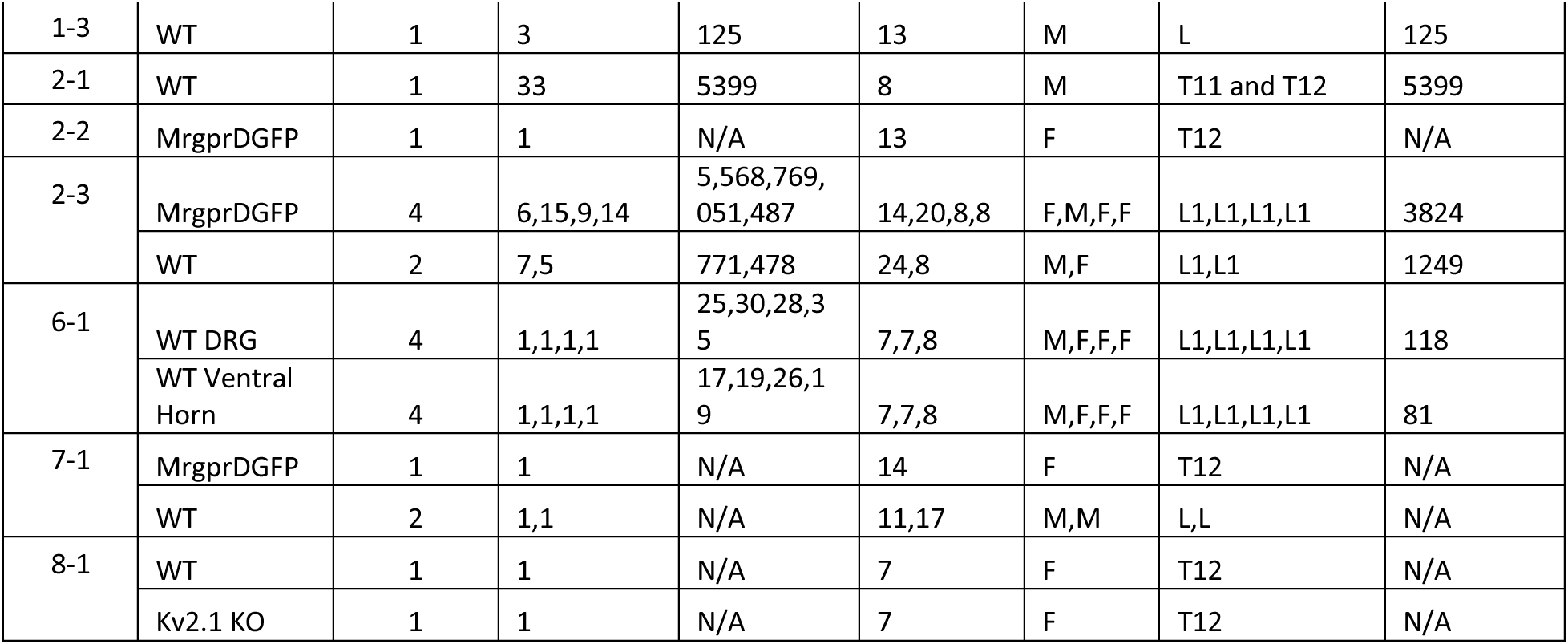
Detailed information on mice used in each figure. In columns “Sections/Plates Analyzed”, “ROIs Analyzed”, “Age”, “Sex” and “Level of Spinal Column” information for each mouse is separated by commas. “ROIs Analyzed” corresponds to ROIs generated by automated analysis. In column “Level of Spinal Column” L1 refers to the first lumbar DRG and T12 refers to the twelfth thoracic DRG and if only L is shown the specific DRG is not known but the DRG is from the lumbar region.

### Human Tissue Collection

Human DRG were obtained in collaboration with Mid-America Transplant (St. Louis, MO) from three donors. Donor #1 was a 59-year-old Caucasian male and the DRG was from the 2^nd^ lumbar region (cause of death: anoxia/stroke). Donor #2 was a 59-year-old black male and the DRG was from the 3^rd^ lumbar region (cause of death: hemorrhagic stroke). Donor #3 was a 58-year-old Caucasian female and the DRG was from the 3^rd^ lumbar region (cause of death: cerebrovascular/stroke). DRG were extracted less than 2 hours after aortic cross clamp and transported to the lab where they were dissected to remove the dura, embedded in Optimal Cutting Temperature (OCT) compound, snap frozen, and stored at -80 °C until use (Valtcheva et al., 2016). Human DRG were obtained from organ donors with full legal consent for use of tissue for research in compliance with procedures approved by Mid-America Transplant.

### Tissue Preparation

Mice were briefly anesthetized with 3-5% isoflurane and then decapitated. The spinal column was dissected, and excess muscle tissue removed. The spinal column was then bisected in the middle of the L1 vertebrae identified by the 13^th^ rib and drop fixed for 1 hour in ice cold PFA: 4% formaldehyde prepared fresh from paraformaldehyde in 0.1 M phosphate buffer (PB) pH adjusted to 7.4 with NaOH.

0.1 M PB buffer was diluted from a 0.4 M PB stock solution that was made by diluting 91.37 g of Na_2_H_2_PO_4_ and 20.98 g of Na_2_H_2_PO_4_ • H_2_O in 2 liters of milliQ water. The relatively short fixation period was chosen because a longer fixation period increased the off target secondary antibody immunofluorescence from mouse isotype secondary antibodies (Supplemental Figure 1). The spine was washed 3× for 10 min each wash in PB and cryoprotected at 4 °C in 30% sucrose in PB for 24 hours. The spine was cut into sections containing two vertebra per sample which were frozen in OCT (Fisher cat#4585) and stored at -80 °C until sectioning. Vertebrae position relative to the 13^th^ rib was recorded for each frozen sample to determine the specific vertebrae position in the spinal cord. Samples were cut into 30 μm sections on a freezing stage sliding microtome and were collected on Colorfrost Plus microscope slides (Fisher cat#12-550-19). Slides were stored at -20 °C or immediately used for multiplex immunofluorescence labeling.

Human DRG were cut into 30 μm sections on a freezing stage sliding microtome and were collected on Colorfrost Plus microscope slides. Sections were briefly thawed to adhere to the slide but were immediately returned to the cryostat kept at -20 °C. Slides were removed from the cryostat and immediately transferred to freshly made 4% PFA pH 7.4 for 10 minutes. Slides were then placed in PB solution for 10 minutes and immediately used for multiplex immunofluorescence labeling. Attempts at immunolabeling Kv2 channels in human DRG tissue that was fixed in 4% PFA for 24 hours were not successful.

### Neuron Cell Culture

Cervical, thoracic and lumbar DRGs were harvested from mice and transferred to Hank’s buffered saline solution (HBSS) (Invitrogen). Ganglia were treated with collagenase (2 mg/ml; Type P, Sigma-Aldrich) in HBSS for 15 min at 37°C followed by 0.05% Trypsin-EDTA (Gibco) for 2.5 min with gentle rotation. Trypsin was neutralized with culture media (MEM, with L-glutamine, Phenol Red, without sodium pyruvate) supplemented with 10% horse serum (heat-inactivated; Gibco), 10 U/ml penicillin, 10 μg/ml streptomycin, MEM vitamin solution (Gibco), and B-27 supplement (Gibco). Serum-containing media was decanted and neurons were triturated using a fire-polished Pasteur pipette in MEM culture media containing the supplements listed above. Neurons were plated on laminin-treated (0.05 mg/ml, Sigma-Aldrich) 35 mm glass bottom dishes (MatTek cat#P35G-1.5-7-C). Neurons were then incubated at 37°C in 5% CO_2_. Neurons were used for imaging experiments within 24 hours after plating.

### Labeling Endogenous Kv2 in live cultured DRG neurons

Plates were removed from 37°C incubator 24 hours after plating. Neurons were washed once with 1 mL neuronal external (NE) solution (3.5 KCl, 155 NaCl, 10 HEPES, 1.5 CaCl2, and 1 MgCl2, 10 mM glucose adjusted to pH 7.4 with NaOH). Neurons were then incubated with NE solution supplemented with 1% BSA for 5 minutes. Neurons were then washed once with NE solution and incubated in 5 µg/mL wheat germ agglutinin conjugated to Alexa Fluor 405 (Biotium cat#29028-1) diluted in NE solution for 2.5 minutes. Neurons were washed twice with NE solution and imaged. NE solution was removed and then neurons were incubated for 10 minutes in 100 nM GxTX-594 prepared as described (Thapa et al., 2021).

Neurons were then washed three times in NE solution and imaged again. The temperature of the imaging chamber was 25 °C throughout the experiment. Experimenter was blinded to which culture dishes contained DRG neurons from either WT or Kv2.1/Kv2.2 double knockout mice throughout the experiment and image analysis.

### Immunohistochemistry

A hydrophobic barrier was drawn around tissue sections mounted on slides as described above using a hydrophobic barrier pen (Scientific Device cat#9804-02). Sections were incubated in 4% milk, 0.1 M phosphate buffer (PB) and 0.2% Triton X-100 (vehicle) for 1 hour. Sections were then incubated in vehicle containing 0.1 mg/mL IgG F(ab) polyclonal IgG antibody (Abcam cat#ab6668) for 1 hour. We determined that a concentration of 0.1 mg/mL maximally blocked off-target secondary antibody labeling when using isotype-specific anti-mouse secondary antibodies (Supplemental Figure 2). Sections were washed 3× for 5 min each in vehicle and then incubated in vehicle containing primary antibodies for 1 hour. Primary antibodies and concentrations used are listed in Table 2. Sections were then washed 3× for 5 min each in vehicle and then incubated in vehicle containing IgG-subclass-specific secondary Abs for one hour (Thermo Fisher). Sections were then washed 3× for 5 min each in PB and mounted with Prolong Gold (Thermo Fisher cat#P36930) and Deckglaser cover glass (Fisher Scientific cat#NC1776158). All incubations and washes were done at room temperature with gentle rocking. Human tissue was immunolabeled identically to mice with the exception that the step incubating sections in vehicle containing 0.1 mg/mL IgG F(ab) polyclonal IgG antibody was omitted and sections were incubated in primary antibodies for 2 hours instead of 1. Primary antibodies used in Figure 2 and Figure 3 were used at saturating concentrations (Supplemental Figure 4).

**Table 2.** Detailed information on antibodies used throughout the manuscript. In column “Concentration Used” if only dilution is given concentration was unknown however, tissue culture supernatant concentrations of antibodies typically range between 15-30 µg/mL.

| Antigen | Antibody name | Species/isotype | Manufaturer information | Concentration used | RRID | Figures |
| --- | --- | --- | --- | --- | --- | --- |
| Kv2.1 | K89/34 | Mouse IgG1 mAb | NeuroMab | 6.7 ug/mL | AB_10672253 | 1, 2, 3, 4,6-1, 7, 8-1, 10, 11, 11-1, 11-2 |
| Kv2.2 | Kv2.2c | Rabbit pAb | In-house Trimmer laboratory | 1:100 purified from strip assay | AB_2801484 | 3, 4, 6-1, 7, 8, 9, 10, 11, 11-1, 11-2,11-3 |
| Nav1.8 | N134/12 | Mouse IgG2a mAb | NeuroMab | 1:5 tissue culture supernatant | AB_10672261 | 1,11-1 |
| Kv2.1 | K89/34R | Mouse IgG2a mAb | In-house Trimmer laboratory | 1:5 tissue culture supernatant | AB_2315768 | 11-1 |
| beta III Tubulin | NA | Rabbit pAb | Abcam ab18207 | 0.33 ug/mL | AB_444319 | 2, 8-1 |
| Kv2.2 | N372B/60 | Mouse IgG2b mAb | NeuroMab | 1 ug/mL | AB_2315868 | 1, 1-3, 6, 10 |
| NF200 | NA | chicken pAb | Abcam ab4680 | 0.2 ug/mL | AB_304560 | 3, 8, 10 |
| Caspr | K65/35 | Mouse IgG1 mAb | NeuroMab | 1:5 tissue culture supernatant | AB_2877274 | 9 |
| Kv1.2 | K14/16 | Mouse IgG2b mAb | NeuroMab | 1:5 tissue culture supernatant | AB_2877295 | 9,11 |
| CGRP | NA | Rabbit pAb | Immunostar | 1:1000 | AB_572217 | 10 |
| DsRed | NA | Rabbit pAb | Takara Cat#632543 | 0.17 ug/mL | AB_2307319 | 10 |
| Nav1.7 | N68/6 | Mouse IgG1 mAb | NeuroMab | 1:5 tissue culture supernatant | AB_2184355 | 11-1,11-3 |

### Imaging

Images were acquired with an inverted scanning confocal and airy disk imaging system (Zeiss LSM 880 Airyscan, 410900-247-075) run by ZEN black v2.1. Laser lines were 405 nm, 488 nm, 543 nm and 633 nm. Low-magnification images to image whole mouse DRG were acquired in confocal or airy disk imaging mode with a 0.8 NA 20x objective (Zeiss 420650-9901) details in figure legends. Images containing both the DRG and spinal cord were tile scan images acquired in confocal mode with the same 20x objective. When imaging whole DRG for Figure 2 and Figure 3 the imaging plane was selected using fluorescence from channels that did not contain anti-Kv2.1 or anti-Kv2.2 immunofluorescence. High-magnification images were acquired in airy disk imaging mode with a 1.4 NA 63x oil objective (Zeiss 420782-9900-799). Linear adjustments to contrast and brightness and average fluorescence intensity z-projections were performed using ImageJ software.

### Image Analysis

Images were analyzed using ImageJ software (Schindelin et al., 2012). A summary of automated analysis for selecting neurons in DRG sections can be found in Supplemental Figure 5. The ImageJ plugin MorphoLibJ was used for performing watershed segmentation of images (Legland, Arganda-Carreras, & Andrey, 2016). Automatic generation of ROIs from watershed segmentation did not distinguish the presence of nuclei and is thus expected to over-count larger diameter neurons. Unless stated otherwise the region of DRG neurons used to analyze anti-Kv2 immunofluorescence is the outer edge (Supplemental Figure 5 I). Immunofluorescence is defined as the raw pixel values from confocal images and was not background subtracted. We noted that in smaller neurons the nucleus takes up a greater percentage of the total volume and could skew fluorescence measurements if fluorescence in the entire soma volume is measured. As Kv2 protein is not present in the nucleus and is enriched at the outer edge (Figure 1) we found that measuring anti-Kv2 immunofluorescence at the outer edge of the neuron reduces potential error that could be associated with soma diameter. For proteins that were not Kv2 channels we analyzed the immunofluorescence of the entire soma as these proteins did not exhibit the same enrichment at the outer edge as Kv2 channels (Figure 9 B). Where specified, manual ROIs were used in analysis otherwise automatically generated ROIs were used. When analyzing immunofluorescence from human neurons, ROIs were drawn around areas within the neuronal soma that do not contain apparent lipofuscin (Supplemental Figure 11 C). Fitting of imaging data was performed using Igor Pro software version 8 (Wavemetrics, Lake Oswego, OR) that employs nonlinear least square curve fitting via the Levenberg-Marquardt algorithm. Distributions of fluorescence intensity from DRG neurons were fit with a log normal distribution:

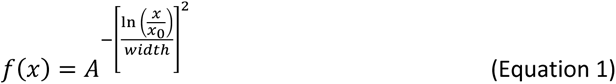

**Figure 1.**
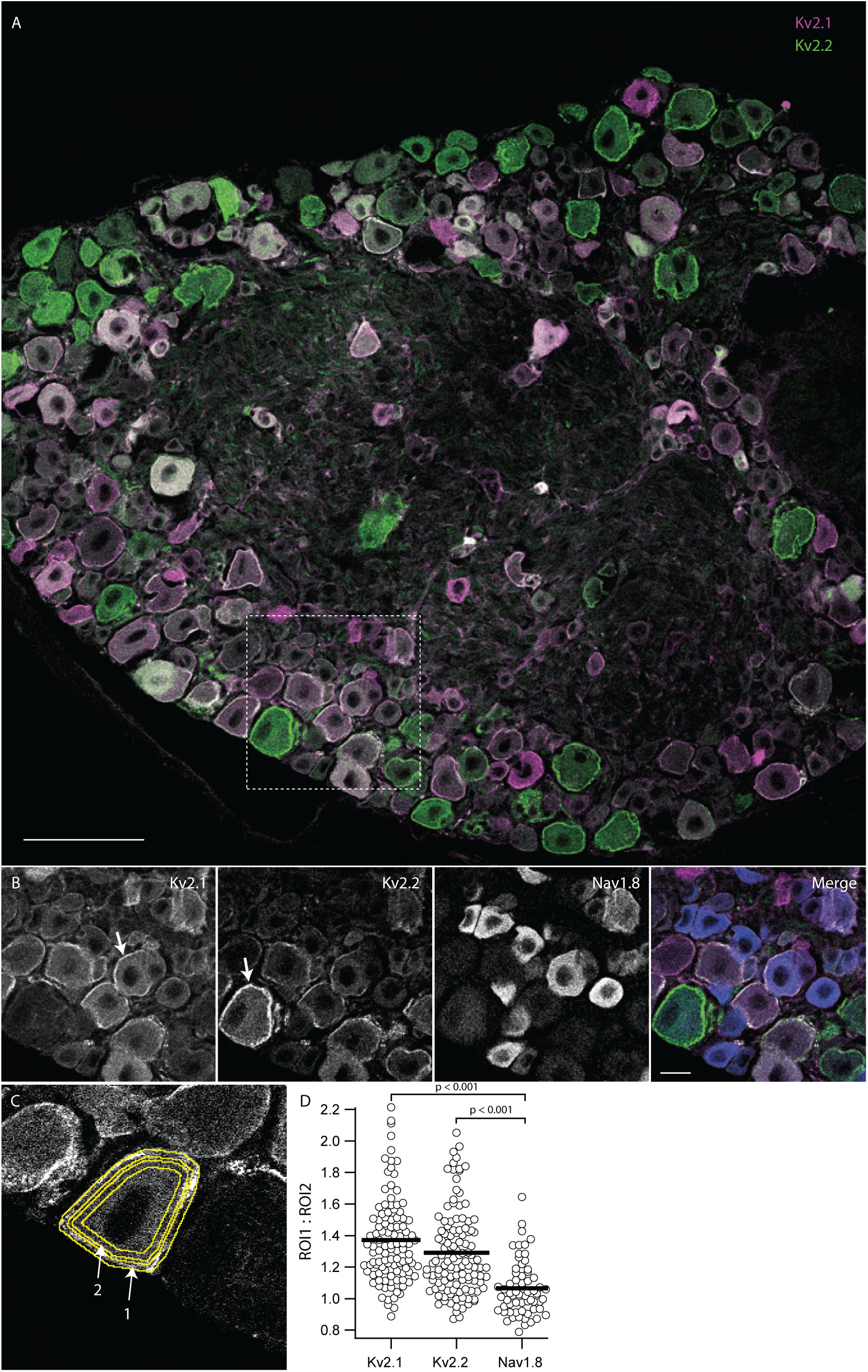
Kv2.1 and Kv2.2 protein are enriched at the outer edge of DRG neuron somas relative to Nav1.8. Lumbar DRG section from an 11 week old male mouse labeled with antibodies which target Kv2.1, Kv2.2 or Nav1.8. ***A***, Anti-Kv2.1 (magenta) and anti-Kv2.2 (green) immunofluorescence in a lumbar DRG section. Scale bar is 100 μm. ***B***, Anti-Kv2.1, anti-Kv2.2 and anti-Nav1.8 immunofluorescence from box shown in A. Arrows indicate prominent localization of anti-Kv2 immunofluorescence at the edge of DRG neuron somas. In merge image anti-Kv2.1, anti-Kv2.2 and anti-Nav1.8 immunofluorescence is magenta, green and blue respectively. Scale bar is 20 μm. ***C***, Representative ROIs that encompass the outer edge of DRG neurons (arrow 1) and the region just inside the outer edge (arrow 2). ***D***, Ratio of anti-Kv2.1, anti-Kv2.2 or anti-Nav1.8 immunofluorescence from outer and inner ROIs for individual neurons from image in A. Bars represent mean. One-way ANOVA p < 0.001. p values in figure represent post hoc Tukey’s test. N = 1 mouse, n = 124 neurons. Detailed information on mouse used can be found in table 1.

Where A = amplitude, x_0_ = mean, and width = √2 times standard deviation. Concentration-effect experiments in Supplemental Figure 4 were fit with the Hill equation:

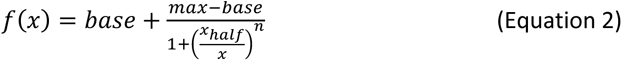

### Code Accessibility

In-house Fiji macros and R script used to process imaging data are available for download at https://github.com/SackLab/DRG-Image-Processing

### Statistics

All statistical tests were performed in Igor Pro software version 8 (Wavemetrics, Lake Oswego, OR). Independent replicates (n) are individual neurons while biological replicates (N) are individual mice. The n and N values for each figure are listed in Table 1. Details of statistical tests are in the figure legends. We did not observe a difference between males and females in detectable Kv2.1 or Kv2.2 protein, but animal numbers were not sufficient for rigorous statistical comparison.

## RESULTS

### Kv2.1 and Kv2.2 immunolabeling is enriched at the apparent plasma membrane of DRG neuron somata

To assess the cellular expression and subcellular localization of Kv2.1 and Kv2.2 voltage gated potassium channels in DRG neurons we used a set of anti-Kv2.1 and anti-Kv2.2 antibodies that have previously been validated in brain tissue from Kv2 knockout mice (Bishop et al., 2015) to perform multiplex immunofluorescence labeling of DRG neurons. Anti-Kv2.1 (magenta) and anti-Kv2.2 (green) immunofluorescence is prominent in DRG neuron somata (Figure 1 A). In neurons we observed anti-Kv2.1 and anti-Kv2.2 immunofluorescence enriched at the outer edge of DRG neuron somata consistent with their plasma membrane localization (Figure 1 B arrows). This putative cell surface localization is distinct from prior reports of anti-Kv2 immunofluorescence in DRG (Tsantoulas et al., 2012) as well as that for other ion channel proteins such as Nav1.8 (Shields et al., 2012) and TRPV1 (Cho & Valtschanoff, 2008), all of which are primarily in the cytoplasm. Anti-Kv2.1 or anti-Kv2.2 immunofluorescence at the edge of neuron somata was more apparent when imaging thinner optical sections. To determine if enrichment of Kv2 immunolabeling at the outer edge of DRG neurons was an artifact of our immunohistochemistry or imaging protocols, we additionally labeled DRG sections with a knockout-validated antibody against Nav1.8. In the same DRG section labeled for Kv2 channels, cytoplasmic anti-Nav1.8 immunofluorescence was prominent in small to medium diameter lumbar DRG neurons, consistent with previous reports (He et al., 2010; Shields et al., 2012) (Figure 1 B). To quantify enrichment of anti-Kv2 versus anti-Nav1.8 immunofluorescence at the outer edge of DRG neurons, we manually drew regions of interest (ROIs) for each DRG soma. We generated a 2 μm wide annulus that encompassed the outer edge and a second 2 μm wide annulus just inside the first one (Figure 1 C arrows 1 and 2 respectively). By comparing the fluorescence intensity of immunofluorescence signals within these annuli we found that the edge of DRG neuron somata is enriched for anti-Kv2.1 and anti-Kv2.2 immunofluorescence relative to that of Nav1.8 (ANOVA < 0.001) (Figure 1 D). Similarly, anti-Kv2.1 and anti-Kv2.2 immunofluorescence is enriched at the edge of DRG neuron somata relative to anti-TRPV1 immunofluorescence (ANOVA < 0.001) (Supplemental Figure 3).

### The majority of dorsal root ganglion neurons express Kv2 protein

To determine whether the anti-Kv2.1 immunofluorescence signal in DRG sections is specific and dependent on the presence of Kv2.1 protein, we compared fluorescence intensities in DRG samples prepared in parallel from age and sex matched wild-type (WT) mice and Kv2.1 knock-out (KO) mice. We observed anti-Kv2.1 immunolabeling in WT DRG neurons that was absent from Kv2.1 KO, while fluorescence corresponding to an antibody that targets βIII tubulin was similar in both WT and Kv2.1 KO mice (Figure 2 A). To quantify fluorescence intensities of DRG sections, we manually drew ROIs around neuron soma profiles with clearly visible nuclei and measured the fluorescence intensity at the outer edge of profiles as shown in Figure 1 C. We define a profile as a slice of a DRG neuron in a histological section (Coggeshall, 1992). We found that 99% of manually identified profiles from the WT mouse DRG shown in Figure 2 A had anti-Kv2.1 immunofluorescence above the mean of Kv2.1 KO mice (Figure 2 B). To reduce human bias in identification of profiles we used an automated method to generate ROIs. As with the manual method, the automated method reliably identified neuronal profiles (Supplemental Figure 5 I). However, the automated method did not distinguish profiles without nuclei which could lead to overrepresentation of larger neurons (Coggeshall, 1992; Coggeshall & Lekan, 1996). Additionally, the automated method occasionally selected ROIs which did not appear to be neurons (Supplemental Figure 5 H red arrows). Despite these limitations, the absolute reproducibility of the automated method when scaled to identify thousands of neuronal profiles indicated that automated ROIs could provide a rigorous means of identifying neuronal profiles for statistical analysis. In automatically generated ROIs from the same WT mouse manually analyzed in Figure 2 B we observed a reduction in the mean anti-Kv2.1 but not anti-βIII immunofluorescence in Kv2.1 KO mice (Figure 2 C and D). We found that 89% of automatically generated ROIs had anti-Kv2.1 immunofluorescence above the mean of the paired Kv2.1 KO mouse. We manually identified that 9% of the automatically generated ROIs from the WT mouse did not appear to be neurons, suggesting that 98% of profiles in this automated dataset could have anti-Kv2.1 immunofluorescence above the mean of the Kv2.1 KO mouse, consistent with manual analysis.

**Figure 2.**
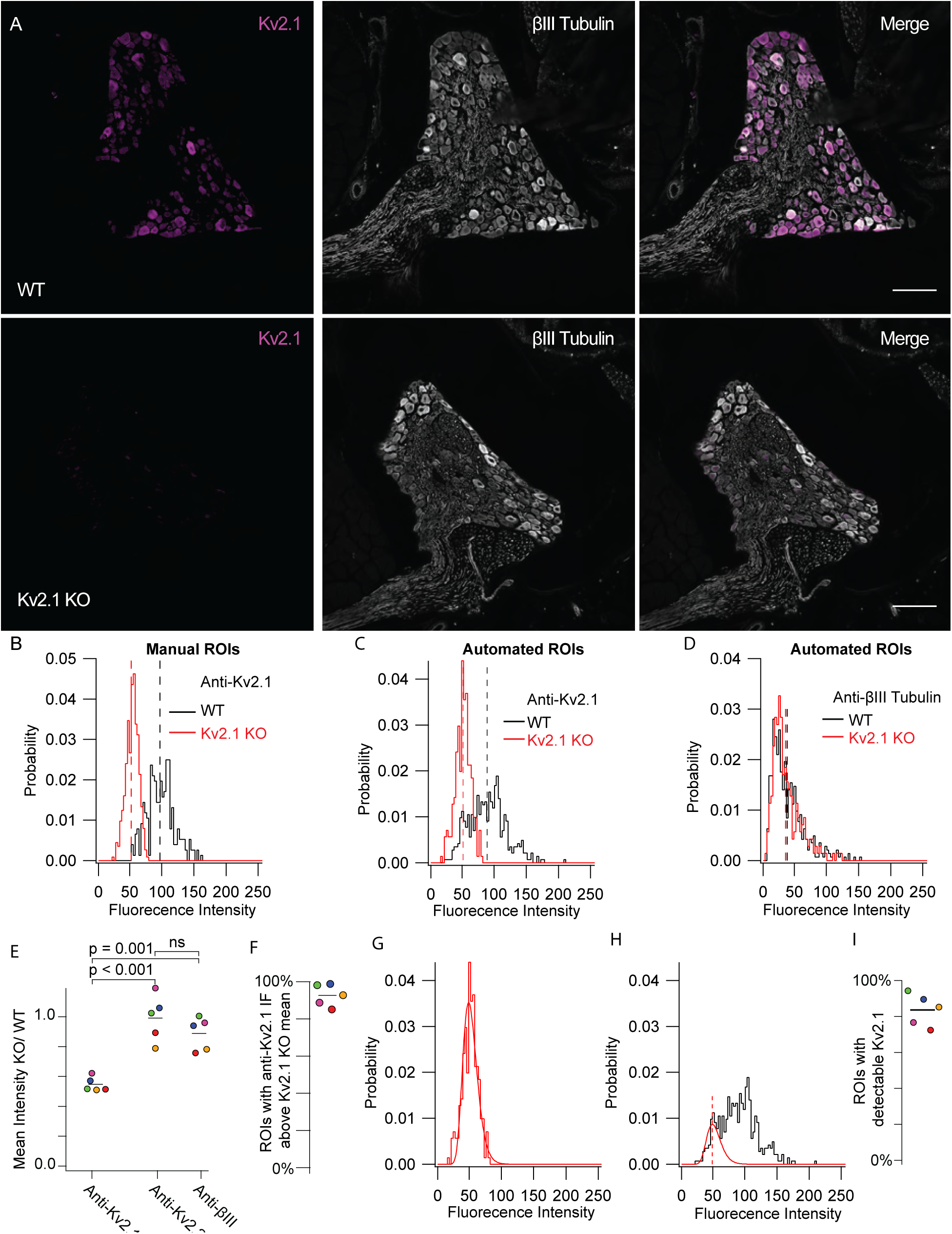
Kv2.1 protein is detectable in mouse DRG neurons. ***A***, WT (top) and Kv2.1 KO (bottom) DRG sections taken from 7 week old female mice from the 13^th^ thoracic DRG immunolabeled for Kv2.1 (magenta) and BIII tubulin (white). Images were taken with identical imaging settings and are set to the same brightness and contrast. Scale bars are 100 µm. ***B***, Distribution of fluorescence intensity from manual analysis of WT (black) and Kv2.1 KO (red) neurons. Dotted lines represent mean. Data represent fluorescence intensity of 254 WT profiles from 10 DRG sections from 1 mouse or 375 Kv2.1 KO profiles from 5 DRG sections from 1 mouse. Images shown in A represent one section from WT and Kv2.1 KO mice used in this data set. ***C***, Distribution of fluorescence intensity from automated analysis of the same data set shown in B. Dotted lines represent mean. Data represent fluorescence intensity of 476 WT or 576 Kv2.1 KO profiles selected by automated analysis method. ***D***, Distribution of BIII tubulin fluorescence intensity from the same WT (black) and Kv2.1 KO (red) profiles shown in C. Dotted lines represent mean. ***E***, Mean fluorescence intensity of Kv2.1 KO neurons normalized to WT neurons labeled with anti-Kv2.1, anti-Kv2.2 and anti-BIII tubulin antibodies. Each point represents one Kv2.1 KO mouse normalized to one age and sex matched WT mouse which was stained simultaneously and imaged with identical microscopy settings. The color of each point represents the same mouse and purple points represent data from the mouse whose DRG immunofluorescence data are shown in A, B, C and D. one-way ANOVA p < 0.001. p values in figure represent post hoc Tukey’s test. ***F***, Percentage of ROIs with anti-Kv2.1 immunofluorescence above the mean immunofluorescence of 5 mice (1 female and 4 male). Point colors correspond to the WT mice analyzed in E. All mice were compared to age and sex matched Kv2.1 KO mice. N = 5 WT and 5 Kv2.1 KO mice. ***G***, Kv2.1 KO data shown in B fit with a log normal distribution (red fit). ***H***, WT data shown in B fit with the Kv2.1 KO distribution (red fit) where width and mean were constrained to the Kv2.1 KO distribution and amplitude was unconstrained (equation 1). Red dotted line represents the mean of the Kv2.1 KO distribution. Only WT data to the left of red dotted line was used for the fit. ***I***, Percentage of ROIs with detectable Kv2.1 protein of 5 mice (1 female and 4 males). Point colors correspond to the WT mice analyzed in E. All mice were compared to age and sex matched Kv2.1 KO mice. N = 5 WT and 5 Kv2.1 KO mice. Detailed information on each mouse used can be found in table 1.

We expanded the automated method to 5 pairs of age and sex matched WT and Kv2.1 KO mice, and found that anti-Kv2.1 immunofluorescence was significantly reduced in each matched Kv2.1 KO mouse relative to anti-Kv2.2 or anti-βIII tubulin immunofluorescence while no significant difference was observed between anti-Kv2.2 and anti-βIII tubulin (Figure 2 E) (ANOVA p < 0.001). We identified that 93 ± 6% (SD) of ROIs from the five WT mice had anti-Kv2.1 immunofluorescence above the mean of age and sex matched Kv2.1 KO mice (Figure 2 F). This analysis confirms that anti-Kv2.1 immunofluorescence reveals Kv2.1 protein at neuron surfaces.

We developed a method to estimate the fraction of ROIs that contain detectable Kv2.1 protein. The histogram of fluorescence intensity in automatically generated ROIs from Kv2.1 KO DRG sections labeled with an anti-Kv2.1 antibody had variability which could reasonably be fit with a log normal distribution (equation 1) (Figure 2 G). We assumed a similar distribution of background immunofluorescence is also present in neuron profiles of age and sex matched WT mice and fit the Kv2.1 KO distribution to the WT histogram to estimate the fraction of ROIs without detectable Kv2.1 (Figure 2 H red gaussian). Using this method, we compared Kv2.1 immunofluorescence in DRG sections of multiple age and sex matched WT and Kv2.1 KO mice. The fitting indicated that 84 ± 9% (SD) of automatically generated ROIs from WT mice have detectable Kv2.1 protein (N = 5 pairs of mice) (Figure 2 I). As approximately 9% of ROIs are expected to not contain neurons these results suggest that greater than 90% of mouse DRG neurons have detectable Kv2.1 protein. We further validated our automated methodology with a transgenic MrgprD-GFP mouse line that expresses GFP in non-peptidergic nociceptors that comprise approximately 19-24% of all lumbar DRG neurons (Dirajlal, Pauers, & Stucky, 2003; Dong, Han, Zylka, Simon, & Anderson, 2001; Wang & Zylka, 2009). In comparing the MrgprD-GFP mice to WT C57BL/6J mice we found that 23 ± 15% (SD) of automatically generated profiles (N = 4 pairs of mice) had detectable GFP (Supplemental Figure 6).

We compared anti-Kv2.2 immunofluorescence from an age and sex matched WT and Kv2.2 KO mouse by drawing ROIs around neuron soma profiles with clearly visible nuclei. This identified that greater than 99% of WT profiles have anti-Kv2.2 immunofluorescence above the mean immunofluorescence of the paired Kv2.2 KO mouse (Figure 3 B). Automatically generated ROIs for multiple age and sex matched WT and Kv2.2 KO mice indicate that 91 ± 5% (SD) of ROIs from WT mice have detectable Kv2.2 protein (N = 8 pairs of mice) (Figure 3 F-H). Using the automated analysis, we found that anti-Kv2.2 immunofluorescence was significantly reduced in Kv2.2 KO mice relative to anti-Kv2.1 or anti-NF200 immunofluorescence in age and sex matched WT and Kv2.2 KO mice (ANOVA p < 0.001) while no significant difference between anti-NF200 or anti-Kv2.1 immunofluorescence was observed (Figure 3 A-E). These combined results suggest that more than 90% of neuron profiles contain detectable Kv2.1 or Kv2.2 protein, raising the possibility that all mouse DRG neurons express Kv2 channels.

**Figure 3.**
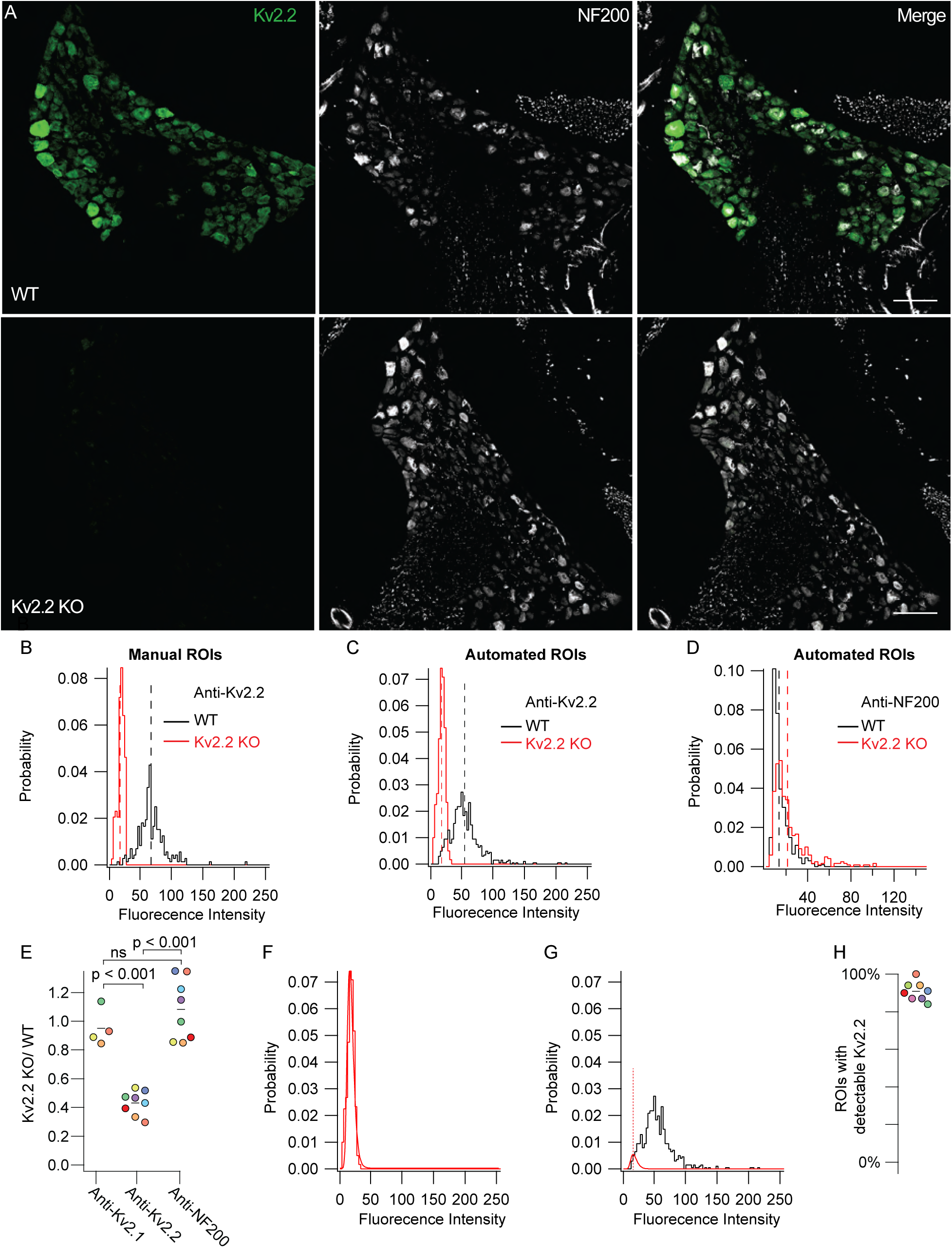
Kv2.2 protein is detectable in mouse DRG neurons. ***A***, WT (top) and Kv2.2 KO (bottom) DRG sections from the 13^th^ thoracic DRG in 7 week old male mice immunolabeled for Kv2.2 (green) and NF200 (white). Identical imaging and display settings. Scale bars are 100 µm. **B**, Distribution of fluorescence intensities from manual analysis of WT (black) and Kv2.2 KO (red) profiles. Dotted lines represent mean. Data represents fluorescence intensities from 241 WT profiles from 11 DRG sections and 1 mouse or 130 Kv2.2 KO profiles from 6 DRG sections and 1 mouse. Images shown in A represent one section from WT and Kv2.1 KO mice used in this data set. ***C***, Distribution of fluorescence intensity from automated analysis of the same data set shown in B. Dotted lines represent mean. Data represent fluorescence intensity of 673 WT or 400 Kv2.2 KO profiles selected by automated analysis method. ***D***, Distribution of anti-NF200 immunofluorescence intensity from the same WT (black) and Kv2.2 KO (red) neurons shown in B. Dotted lines represent mean. ***E***, Mean fluorescence intensity of Kv2.2 KO ROIs normalized to WT neurons labeled with anti-Kv2.1, anti-Kv2.2 and anti-NF200 antibodies. Each point represents one Kv2.2 KO mouse normalized to one age and sex matched WT mouse which was stained simultaneously and imaged with identical microscopy settings. The color of each point represents the same mouse and purple points represent data from the male mouse whose DRG immunofluorescence data are shown in A, B and C. Missing points in anti-Kv2.1 column are because some sections were not labeled with anti-Kv2.1 antibodies. one-way ANOVA p < 0.001. p values in figure represent Tukey’s post hoc test. ***F***, Kv2.2 KO data shown in B fit with a log normal distribution (red fit). ***G***, WT data shown in B fit with the Kv2.2 KO distribution (red fit) where width and mean were constrained to the Kv2.2 KO distribution and amplitude was unconstrained (equation 1). Red dotted line represents the mean of the Kv2.2 KO distribution. Only WT data to the left of red dotted line was used for the fit. ***H***, Percentage of ROIs with detectable Kv2.2 protein of 8 mice (7 males and 1 female). Point colors correspond to the WT mice analyzed in D. All mice were compared to age and sex matched Kv2.2 KO mice. N = 8 WT and 8 Kv2.2 KO mice. Detailed information on each mouse used can be found in table 1.

### Kv2.1 immunolabeling decreases in older mice while Kv2.2 immunolabeling remains similar

Kv2.2 transcript in mouse DRG decreases during postnatal development (Regnier et al., 2016) and Kv2.1 protein expression in mouse brain decreases with age (Cotella et al., 2012; Regnier et al., 2016). To identify if Kv2 protein levels in DRG neurons vary between young adult and old mice we compared anti-Kv2.1 and anti-Kv2.2 immunofluorescence from samples subjected to identical immunolabeling and imaging protocols. We observed lower anti-Kv2.1 (Figure 4 A, B and C) in DRG sections from 50 week old mice relative to 7-16 week old mice. Comparisons with Kv2.1 KO mice indicate that 50 ± 23% (SD) of ROIs from 50 week old mice express detectable Kv2.1 protein, significantly less (*p* = 0.018) than the 84 ± 9% (SD) of ROIs with detectable Kv2.1 protein from 7-16 week old mice (Figure 4 D). In contrast, similar anti-Kv2.2 immunofluorescence was seen in samples from young adult and old mice (Figure 4 E, F and G). Comparisons with Kv2.2 KO mice indicate that 96 ± 3% of profiles from 50 week old mice express detectable Kv2.2 protein, a weakly significant increase (*p* = 0.03) from the 88 ± 3% (SD) of profiles with detectable Kv2.2 protein from 7-24 week old mice (Figure 4 H). We note that this weak statistical increase is underpowered due to the availability of only three 50 week old Kv2.2 KO mice.

**Figure 4.**
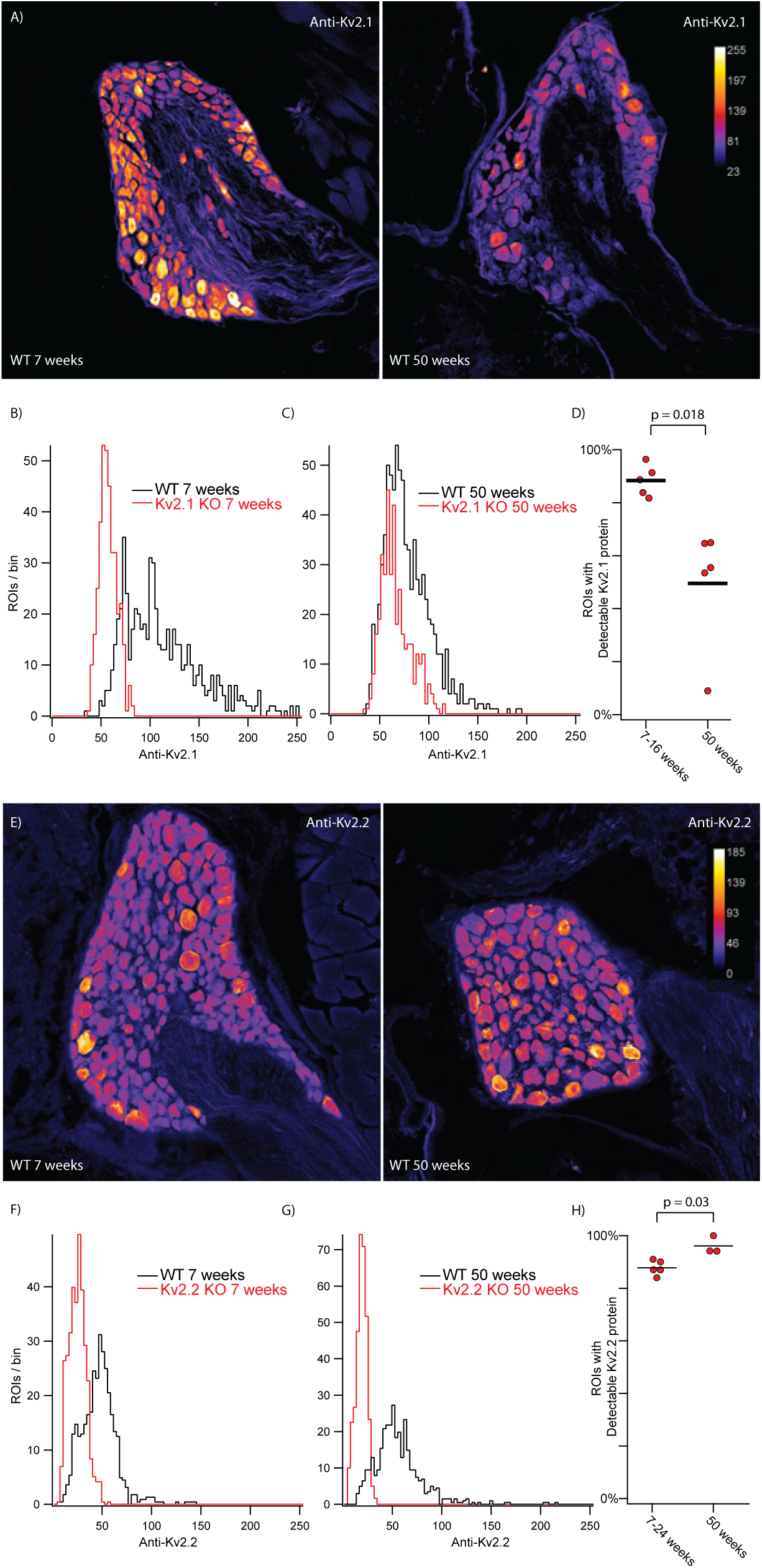
Detectable Kv2.1 protein decreases in older mice while detectable Kv2.2 does not. ***A***, DRG sections from the 13^th^ thoracic DRG in 7 week (left) and 50 week (right) old mice immunolabeled for Kv2.1. Vertical bar on right is pseudo coloring key for pixel intensity. Identical imaging and display settings. Scale bars are 100 µm. ***B***, Distribution of fluorescence intensities from 7 week old WT (black) and 7 week old Kv2.1 KO (red) ROIs generated by automated method. 609 WT ROIs from 1 mouse. 367 Kv2.1 KO ROIs from 1 mouse. ***C***, Distribution of fluorescence intensities from 50 week old WT (black) and 50 week old Kv2.1 KO (red) ROIs. 793 WT ROIs from 1 mouse. 378 Kv2.1 KO ROIs from 1 mouse. ***D***, Percentage of ROIs with detectable Kv2.1 protein in 7-16 week old and 50 week old mice. Data from 7-16 week old mice is the same data in Figure 2. N = 4 mice 7 weeks old and 1 mouse 16 weeks old and N = 5 mice 50 weeks old. Detailed information on each mouse used can be found in table 1. ***E***, DRG sections from the 13th thoracic DRG in 7 week (left) and 50 week (right) old mice immunolabeled for Kv2.2. Vertical bar on right is pseudo coloring key for pixel intensity. Identical imaging and display settings. Scale bars are 100 µm. ***F***, Distribution of fluorescence intensities from 7 week old WT (black) and 7 week old Kv2.2 KO (red) ROIs generated by automated method. 746 WT ROIs from 1 mouse. 717 Kv2.2 KO ROIs from 1 mouse. Data from same mice shown in E. ***G***, Distribution of fluorescence intensities from 50 week old WT (black) and 50 week old Kv2.2 KO (red) ROIs generated by automated method. 671 WT ROIs from 1 mouse. 398 Kv2.2 KO ROIs from 1 mouse. Data from same mice shown in E. ***H***, Percentage of ROIs with detectable Kv2.2 protein in 7-24 week old and 50 week old mice. Data is the same data from Figure 3 where mice were separated into a young group (7-24 weeks) and an old group (50 weeks). N = 3 mice 7 weeks old, 1 mouse 24 weeks old and 1 mouse 25 weeks old. N = 3 mice 50 weeks old. Detailed information on each mouse used can be found in table 1.

### Kv2 channels are expressed on the cell surface of acutely dissociated DRG neurons

To determine if Kv2 channels are present on neuron surfaces we labeled live acutely dissociated DRG neurons with a cell-impermeant fluorescent probe which binds Kv2 channels. The probe is a variant of the tarantula peptide guangxitoxin-1E conjugated to Alex Fluor 594 (GxTX-594) which binds to an extracellular site of Kv2.1 and Kv2.2 channels (Thapa et al., 2021). We applied GxTX-594 to live dissociated DRG neurons from WT mice and double knockout (DKO) mice lacking expression of both Kv2.1 and Kv2.2 protein (Figure 5). In WT DRG neurons, we observed fluorescence at the membrane after application of 100 nM GxTX-594 that was not present before the addition of GxTX-594 (Figure 5 A). Wheat germ agglutinin conjugated to Alexa 405 (WGA-405) was used to identify the surface membrane of cultured DRG neurons and confirmed that the GxTX-594 fluorescence observed in WT neurons was cell surface localized (Figure 5 A middle). The cell surface localized fluorescence of GxTX-594 observed in WT neurons was not present in Kv2.1/Kv2.2 DKO neurons (Figure 5 B). To quantify the cell surface-associated fluorescence signals of individual WT and Kv2.1/Kv2.2 DKO neurons, we used an automated method to generate ROIs corresponding to the WGA-405 fluorescence and measured GxTX-594 labeling within this ROI (Figure 5 C). This analysis confirmed a reduction in GxTX-594 fluorescence at the membrane in cultured DRG neurons from Kv2.1/Kv2.2 DKO mice compared to WT mice (Figure 5 D).

**Figure 5.**
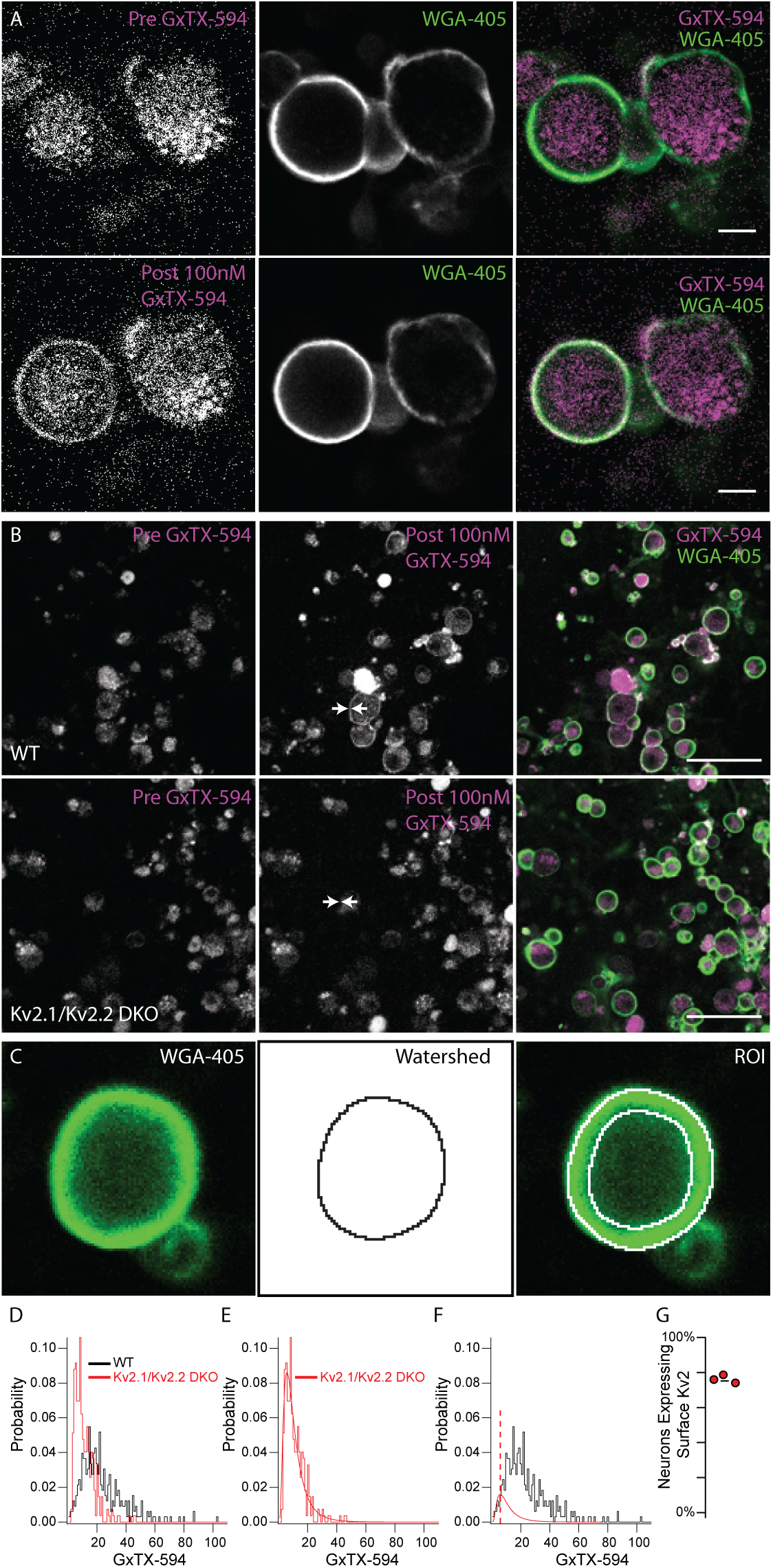
Kv2 channels are expressed at the surface membrane of DRG neurons. ***A***, Fluorescence of live dissociated DRG neurons excited at 594 nm before (top left) and after (bottom left) application of 100 nM GxTX-594. Fluorescence from membrane marker WGA-405 before (top middle) and after application of 100 nM GxTX-594 (bottom middle). Merge image shows 594 excitation fluorescence (magenta) and 405 excitation fluorescence (green). Scale bar 20 µm. ***B***, Dissociated DRG neurons from WT (top) and Kv2.1/Kv2.2 DKO (bottom) mice before and after the application of GxTX-594, left panel and middle panel respectively. Arrows in middle panel indicate location of surface membrane based on WGA-405 fluorescence. Right images are merge of 594 excitation fluorescence (magenta) and 405 excitation fluorescence (green) after application of 100 nM GxTX-594. Scale bars 100 µm. ***C***, Example of WGA-405 fluorescence (left) used in watershed segmentation (middle) to generate annulus ROI (right) used to analyze fluorescence intensity at the membrane. ***D***, Distribution of fluorescence intensity from WT (black) and Kv2.1/Kv2.2 DKO (red) neurons. Data represents the fluorescence intensity of 326 WT neurons from 1 mouse or 271 Kv2.1/Kv2.2 DKO neurons from 1 mouse. DRG from all levels of the spinal cord were pooled. ***E***, Kv2.1/Kv2.2 DKO data shown in D fit with a log normal distribution (red fit). ***F***, WT data shown in D fit with the Kv2.1/Kv2.2 DKO distribution (red fit) where width and mean were constrained to the Kv2.1/Kv2.2 DKO distribution and amplitude was unconstrained (equation 1). Red dotted line represents the mean of the Kv2.1/Kv2.2 DKO distribution. Only WT data to the left of red dotted line was used for the fit. ***G***, Percentage of neurons with detectable surface Kv2 protein, from an experiment where one WT mouse was compared to one DKO mouse and an identical experiment where two WT mice were compared to one DKO mouse (N=3 WT mice N=2 Kv2.1/Kv2.2 DKO mice). N = 3 WT mice and N = 2 DKO mice. Detailed information on each mouse used can be found in table 1.

Similar to the Kv2 immunofluorescence signals from KO mice (Figures 2 and 3), GxTX-594 fluorescence from Kv2.1 /Kv2.2 DKO mice could be reasonably fit with a log normal distribution (equation 1) (Figure 5 E). Applying the same method used to estimate the fraction of neuronal profiles expressing detectable Kv2.1 and Kv2.2 immunofluorescence we compared cultured DRG neurons of age and sex matched WT and Kv2.1/Kv2.2 DKO mice and estimated that 76 ± 2.4% (SD) of DRG neurons have detectable cell surface GxTX-594 fluorescence (Figure 5 F and G). This indicates that at least 76% of neurons express Kv2 channels in their surface membrane, consistent with a report of Kv2 conductance in every DRG neuron type assessed (Zheng et al., 2019).

### Anti-Kv2.2 immunofluorescence is enriched in DRG neurons relative to spinal cord neurons

In mammals, anti-Kv2.1 and anti-Kv2.2 immunofluorescence is apparent in central neurons of the brain (Kihira et al., 2010; Bishop et al., 2015) and anti-Kv2.1 in the spinal cord (Bishop et al., 2015; Muennich & Fyffe, 2004). We measured Kv2 immunofluorescence intensities in sections of mouse spinal column which contain both the DRG and the spinal cord. Neurons in the ventral and dorsal horn of the spinal cord exhibit anti-Kv2.1 immunofluorescence (Figure 6 A). We observed anti-Kv2.1 puncta on neurons in the ventral horn (Figure 6 A inset) similar to that described in alpha motor neurons (Fletcher et al., 2017; Romer et al., 2014). Fewer spinal cord neurons exhibit anti-Kv2.2 immunofluorescence than anti-Kv2.1 (Figure 6 A and Supplemental Figure 7 A arrow heads). Kv2.2 channel protein in the spinal cord of mice has to our knowledge not been assessed, but anti-Kv2.2 immunofluorescence has been observed early in development in the ventrolateral spinal cord in the frog *Xenopus laevis* (Gravagna, Knoeckel, Taylor, Hultgren, & Ribera, 2008). We analyzed anti-Kv2 immunofluorescence from individual neurons in the DRG and ventral horn of five mice and found that while anti-Kv2.1 immunofluorescence is similar between the two regions (Figure 6 B), anti-Kv2.2 immunofluorescence is significantly higher in DRG neurons relative to ventral horn neurons (Figure 6 C). Approximately 30% of neurons in the DRG have anti-Kv2.2 immunofluorescence at a level higher than the brightest anti-Kv2.2 immunofluorescence observed in the spinal cord. We repeated this analysis with a different anti-Kv2.2 antibody and found similar results (Supplemental Figure 7).

**Figure 6.**
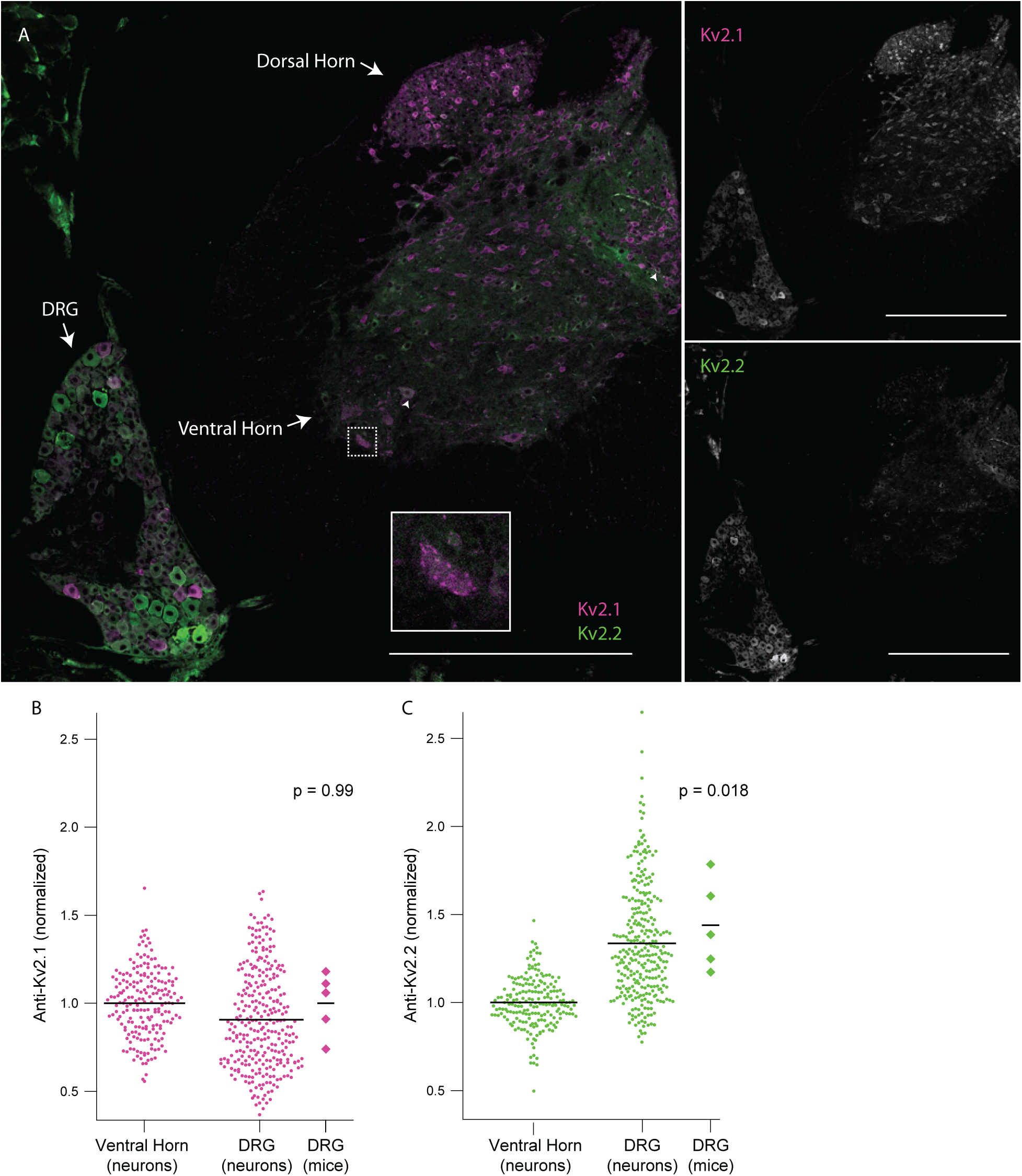
DRG neurons have enriched Kv2.2 protein compared to neurons in the spinal cord. ***A***, Anti-Kv2.1 (magenta) and anti-Kv2.2 (green) immunofluorescence in a spinal cord section from the 2^nd^ lumbar vertebra (left). Anti-Kv2.1 immunofluorescence (right top) and anti-Kv2.2 immunofluorescence (right bottom). Arrows show neurons in the spinal cord with punctate anti-Kv2.1 immunofluorescence. Arrow heads show neurons in the spinal cord with anti-Kv2.2 immunofluorescence. Scale bars are 500 μm. ***B***, Anti-Kv2.1 immunofluorescence from individual neuron profiles (circles) from multiple mice in the DRG and ventral horn normalized to the average fluorescence intensity of neuron profiles in the ventral horn. Diamonds to the right of data represent the average intensity of individual mice. Significant differences from 1 were calculated for individual mice using Students t-test. N = 5 mice, n = 295 in DRG and n = 200 in ventral horn. Detailed information on each mouse used can be found in table 1. ***C***, Identical analysis as in panel B but with anti-Kv2.2 immunofluorescence.

### Kv2 channels on DRG neuron somata and stem axons form clustered subcellular patterns with similarities to and distinctions from central neurons

In neurons throughout the brain (Trimmer, 1991, Scannevin et al., 1996, Bishop et al., 2015) and motor neurons in the spinal cord (Muennich and Fyffe, 2004), Kv2 channels form punctate structures referred to as clusters. In central neurons, Kv2 clusters localize to the soma, proximal dendrites and axon initial segment (Lim, Antonucci, Scannevin, & Trimmer, 2000; Scannevin et al., 1996; Trimmer, 1991). At clusters, Kv2 channels organize protein signaling complexes and endoplasmic reticulum-plasma membrane junctions (Johnson et al., 2018; Kirmiz et al., 2018; Panzera et al., 2022; Vierra et al., 2021). To determine whether Kv2 channels in DRG neurons form clusters similar to central neurons, we compared *en face* z-projections of Kv2 clusters in DRG neurons and ventral horn neurons in the same spinal cord section (Figure 7). Anti-Kv2.1 immunofluorescence in the ventral horn neurons of these mice resembles previous reports of mice (Fletcher et al., 2017) and rats (Fletcher et al., 2017; Muennich & Fyffe, 2004). In both DRG and spinal cord neurons, Kv2.1 and Kv2.2 channels form dense clusters (Figure 7 A and B). Kv2 channel clusters in DRG typically appeared smaller than Kv2 clusters in spinal cord.

**Figure 7.**
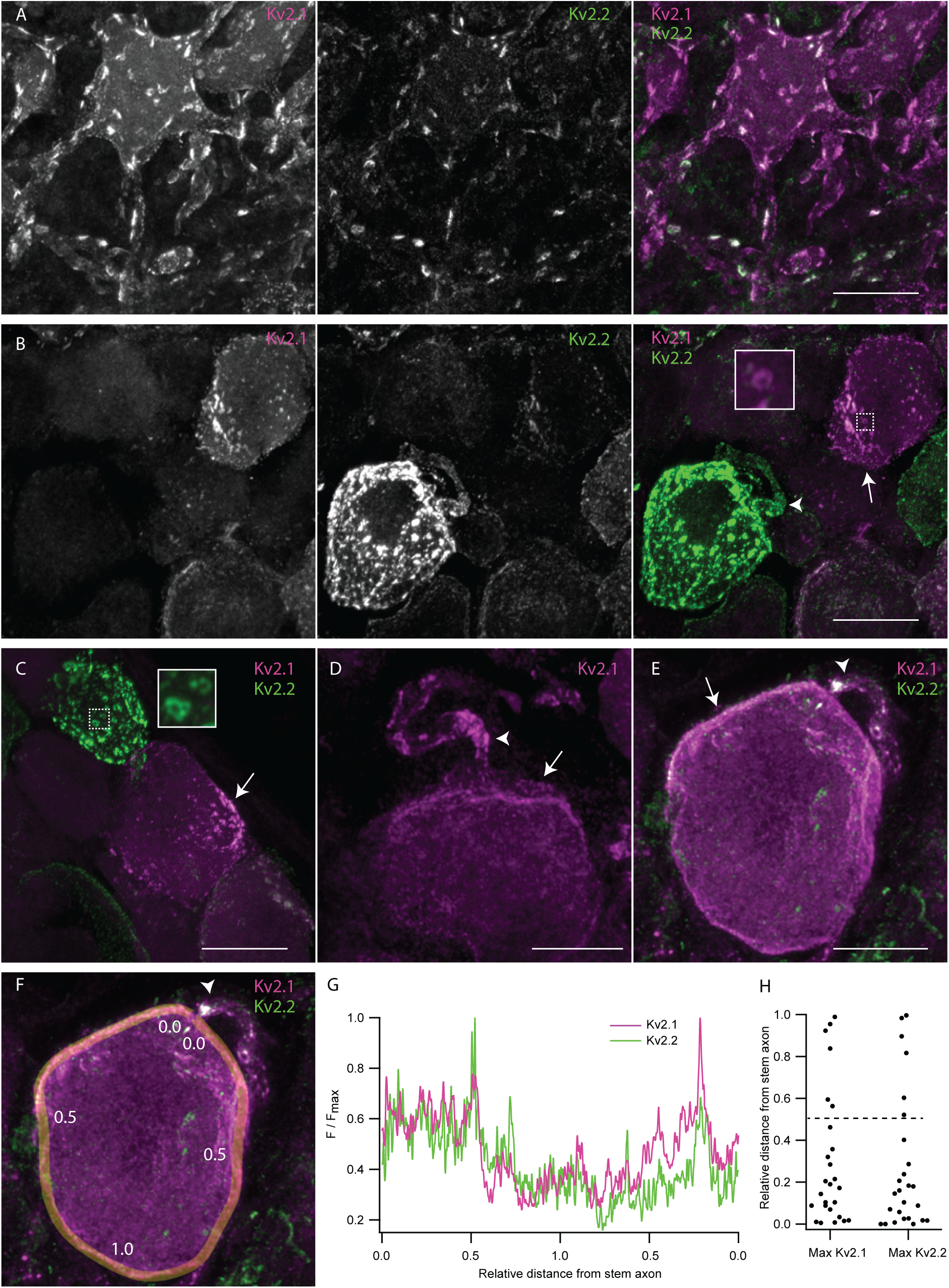
Kv2 channels form clusters on DRG neuron somas and stem axons that are distinct from Kv2 channel clusters on ventral horn neurons. ***A***, Z-projection of anti-Kv2.1 and anti-Kv2.2 immunofluorescence in a ventral horn neuron from the 1^st^ lumbar vertebra of a 7 week old male mouse. ***B***, Z-projection of anti-Kv2.1 and anti-Kv2.2 immunofluorescence in DRG neurons from the same mouse and section of the spinal column as neuron shown in A. Inset is enlargement of the Kv2.1 donut cluster in dotted box. ***C***, Z-projection of anti-Kv2.1 and anti-Kv2.2 immunofluorescence in DRG neurons from the same mouse as A and B. Inset is enlargement of the Kv2.2 donut cluster in dotted box. ***D***, Z-projection of anti-Kv2.1 immunofluorescence in DRG neuron from the 13^th^ thoracic DRG of a 24 week old male mouse. ***E***, Z-projection of anti-Kv2.1 and anti-Kv2.2 immunofluorescence in DRG neuron from same mouse in D. ***F***, Exemplar ROI for analyzing localization of Kv2 channel density relative to stem axon. Same image as E. ROI line width is 1.24 µm. Numbers along line indicate approximate distance from stem axon normalized to the midpoint of the line. ***G***, Anti-Kv2.1 (magenta) and anti-Kv2.2 (green) immunofluorescence intensity along the ROI shown in F. Distance along the line was normalized such that the stem axon is 0 and the midpoint of the line is 1. ***H***, Analysis of the relative distance from the stem axon of the max anti-Kv2.1 or anti-Kv2.2 immunofluorescence. Dotted line represents the middle of neurons relative to the stem axon. In all images arrows indicate asymmetrical clusters of Kv2 channels while arrow heads indicate the apparent stem axons. Display settings are not identical between images. Scale bars are 20 µm.

However, we are not certain these observations in fixed tissue represent the in vivo arrangement, as Kv2.1 channels in brain neurons can decluster in the period between sacrifice and fixation (Misonou et al., 2005). In some DRG neurons, we observed donut shaped Kv2.1 and Kv2.2 clusters (Figure 7 B and C insets) which resemble donut shaped Kv2 clusters in interneurons which harbor specialized protein machinery in their center (Vullhorst et al., 2015).

DRG neurons are distinct from central neurons as they have a single stem axon that exits the cell soma and bifurcates into the central and peripheral axon branches (Ha, 1970). We observed punctate immunofluorescence for both Kv2.1 and Kv2.2 on the apparent stem axons of DRG neurons (Figure 7 B, D and E arrow heads, Supplemental Figure 8). In DRG sections from the *MrgprD^GFP^*mouse line we observed Kv2 clusters on the stem axon of GFP-expressing neurons. As *MrgprD^GFP^* marks non-peptidergic nociceptors (Zheng et al., 2019) we confirm stem axon localization in this subpopulation of unmyelinated neurons (Supplemental Figure 8 A).

While Kv2 clusters in central neurons and many DRG neurons are distributed evenly throughout the soma, we found that DRG neurons asymmetrically distribute Kv2 clusters on the soma (Figure 7 B-E arrows). To test if enrichment of Kv2 channel clusters is oriented to a specific region on the soma we manually traced the surface of neurons whose apparent stem axons were visible (Figure 7 F). The brightest anti-Kv2.1 and anti-Kv2.2 immunofluorescence is near the stem axon in 76 and 80% of DRG neurons respectively (Figure 7 G and H). Asymmetric distribution of Kv2 channels is not frequently observed in central neurons of mice (Bishop et al., 2018; Bishop et al., 2015; Romer et al., 2014) suggesting a potentially unique role of Kv2 subcellular organization in DRG neurons.

### Kv2.2 channels are expressed in peripheral axons of DRG neurons

To determine if anti-Kv2 immunofluorescence is detectible in DRG axons in regions beyond the stem axon we immunolabeled samples containing both peripheral and central axons, using NF200 or βIII tubulin as markers for DRG axons which target myelinated neurons or all DRG neurons respectively. Neurons were not co-labeled with βIII tubulin and Kv2.2 targeting antibodies because they were of the same isotype. We did not observe anti-Kv2.1 immunofluorescence in peripheral axons (Supplemental Figure 9) from the same DRG section that had detectable anti-Kv2.1 immunofluorescence in neuron somas (Figure 2). However, anti-Kv2.2 immunofluorescence is detectable in peripheral and central axons of WT but not Kv2.2 KO mice (Figure 8 B and C). Anti-Kv2.2 immunofluorescence on myelinated peripheral axons appears as discrete puncta or as larger bands which span the width of the axon (Figure 8 B arrows). We also observed anti-Kv2.2 immunofluorescence not colocalized with NF200, suggestive of unmyelinated fibers (Figure 8 B arrow head). In *MrgprD^GFP^*mice, anti-Kv2.2 immunofluorescence colocalizes with axons expressing cytosolic GFP, identifying that non-peptidergic nociceptors also express Kv2.2 protein along the peripheral axon (Figure 8 D). We further investigated the expression of Kv2.2 in peripheral myelinated fibers by co-labeling peripheral axons with antibodies specific for CASPR (Einheber et al., 1997) and Kv1.2 (Rasband et al., 1998) which identify the paranodal and juxtaparanodal regions of nodes of Ranvier, respectively in myelinated fibers. We found punctate anti-Kv2.2 immunofluorescence associated with both CASPR and Kv1.2 in some WT DRG neuron axons that is not present in Kv2.2 KO mice (Supplemental Figure 10 A). We analyzed anti-Kv2.2 immunofluorescence in WT and Kv2.2 KO mice by manually drawing ROIs around CASPR and Kv1.2 immunofluorescence while blinded to anti-Kv2.2 immunofluorescence and found that 26/34 CASPR and 20/31 Kv1.2 immunolabeled regions had anti-Kv2.2 immunofluorescence brighter than the brightest anti-Kv2.2 immunofluorescence measured in Kv2.2 KO mice (Supplemental Figure 10 B). These results indicate that Kv2.2 but not Kv2.1 protein is detectable in axons of myelinated and unmyelinated somatosensory neurons. However, these results do not identify if Kv2.2 channels are localized to specific subcellular regions of somatosensory axons.

**Figure 8.**
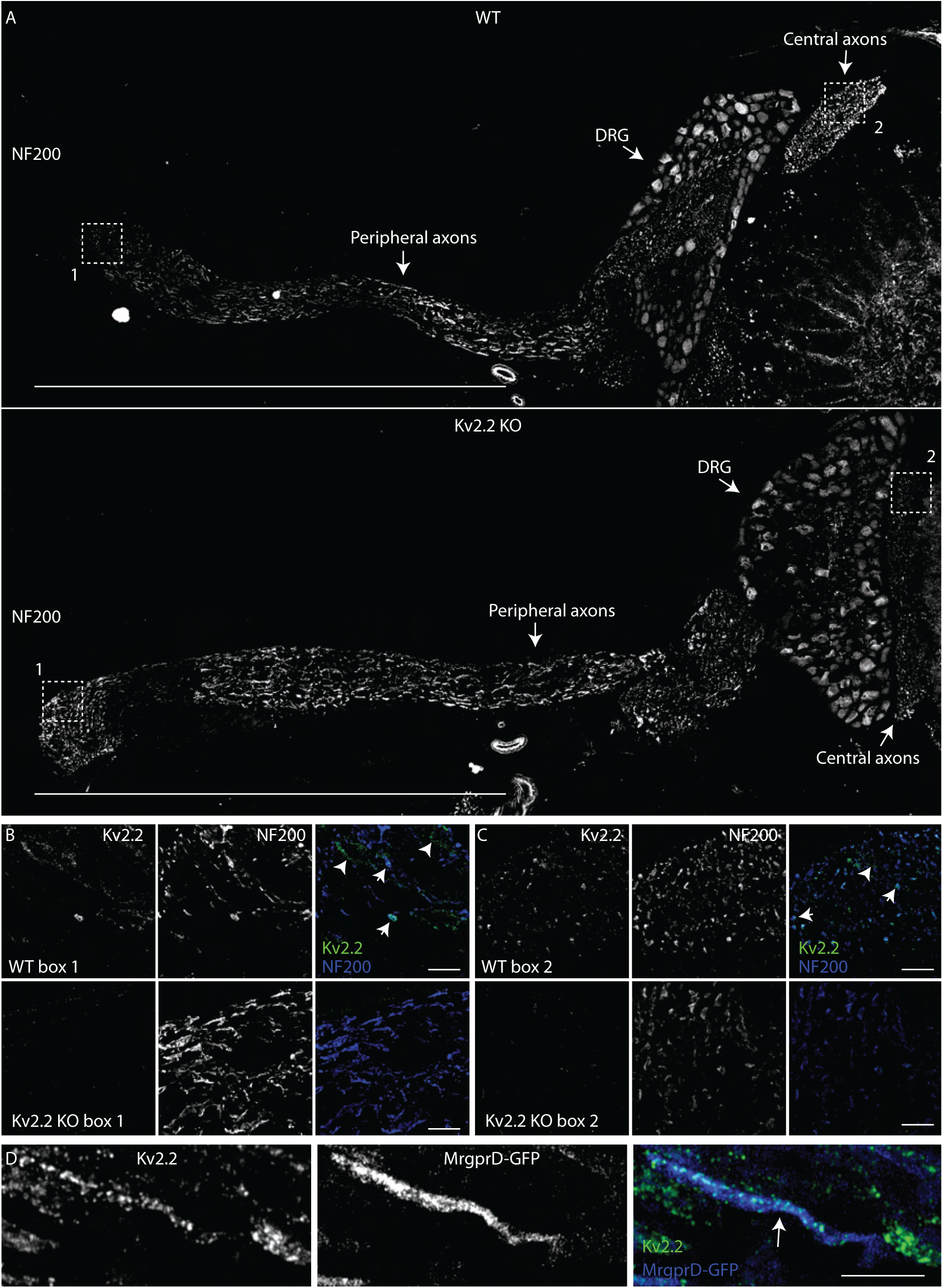
Kv2.2 channels are expressed in peripheral axons of DRG neurons. ***A***, WT (top) and Kv2.2 KO (bottom) sections containing the DRG, peripheral and central axons from the 1^st^ lumbar DRG in age and sex matched 7 week old mice immunolabeled for NF200 (white). Scale bar is 1 mm. ***B***, High magnification z-projection of anti-Kv2.2 and anti-NF200 immunofluorescence from box 1 in A of WT and Kv2.2 KO mice. Arrows indicate myelinated axons which show prominent anti-Kv2.2 immunofluorescence. Scale bars are 20 μm. ***C***, High magnification z-projection of anti-Kv2.2 and anti-NF200 immunofluorescence from box 2 in A of WT and Kv2.2 KO mice. Arrows indicate myelinated axons which show prominent anti-Kv2.2 immunofluorescence. Scale bars are 20 μm. ***D***, High magnification z-projection of anti-Kv2.2 immunofluorescence and MrgprD-GFP fluorescence in the peripheral axons of the 12^th^ thoracic DRG of a 13 week old MrgprD-GFP mouse. Arrow indicates anti-Kv2.2 immunofluorescence on a GFP^+^ axon. Scale bar is 10 μm.

### Anti-Kv2.1 and anti-Kv2.2 immunofluorescence is non-uniform across DRG neuron subtypes

Different voltage gated potassium channels make distinct contributions to outward potassium currents in sensory neuron subtypes (Zheng et al., 2019). We observed subpopulations of DRG neurons that have bright anti-Kv2.1 immunofluorescence but dim anti-Kv2.2 immunofluorescence and vice-versa (Figure 9 A and C, Figure 1 A, and Figure 6 A). We hypothesized that differences in Kv2.1 and Kv2.2 protein density could potentially denote specific subtypes of sensory neurons. In order to test if relative density of Kv2.1 and Kv2.2 protein differs in subtypes of DRG neurons, we assessed anti-Kv2.1 and anti-Kv2.2 immunofluorescence in four DRG neuron subtypes. Neurons identified by genetically encoded markers were GFP-expressing neurons from *MrgprD^GFP^*mice which we refer to as non-peptidergic nociceptors (Zheng et al., 2019), or tdTomato-expressing neurons from *PV^Ai14^* mice which we refer to as proprioceptors (Zheng et al., 2019). Neurons immunolabeled for CGRP we refer to as peptidergic nociceptors (Zheng et al., 2019) and neurons immunolabeled for NF200 we refer to as myelinated DRG neurons (Usoskin et al., 2015) (Figure 9 B and D). We quantified detectable Kv2.1 or Kv2.2 protein density in individual ROIs by rank ordering the immunofluorescence intensity. Rank orders from multiple mice were pooled with a value of 100% representing the brightest ROI in each mouse (Figure 9 D). The rank percentiles of Kv2.1 and Kv2.2 have similar distributions in peptidergic and non-peptidergic nociceptors. Many ROIs with the highest Kv2.1 or Kv2.2 rank percentile were NF200-positive myelinated neurons. Proprioceptors, a subset of the myelinated neurons, appear to account for the subpopulation of neurons with bright anti-Kv2.2 and dim anti-Kv2.1, and have high rank percentiles of Kv2.2 but low rank percentiles of Kv2.1. (Figure 9 A, B, and C). Profiles with bright anti-Kv2.1 and dim anti-Kv2.2 immunofluorescence are mostly NF200-positive myelinated neurons of unknown subtype (Figure 9 A, B, and C blue points). Overall, these results identify that the relative densities of Kv2.1 and Kv2.2 protein are similar in DRG neuron subtypes except proprioceptors. These results are consistent with reports that Kv2.1 and Kv2.2 mRNA transcript levels are similar in DRG neuron subtypes except proprioceptors. Our finding that Kv2.1 protein density is anomalously low in proprioceptors mirrors the finding that Kv2.1 transcripts are also low in proprioceptors (Usoskin et al., 2015; Zheng et al., 2019).

**Figure 9.**
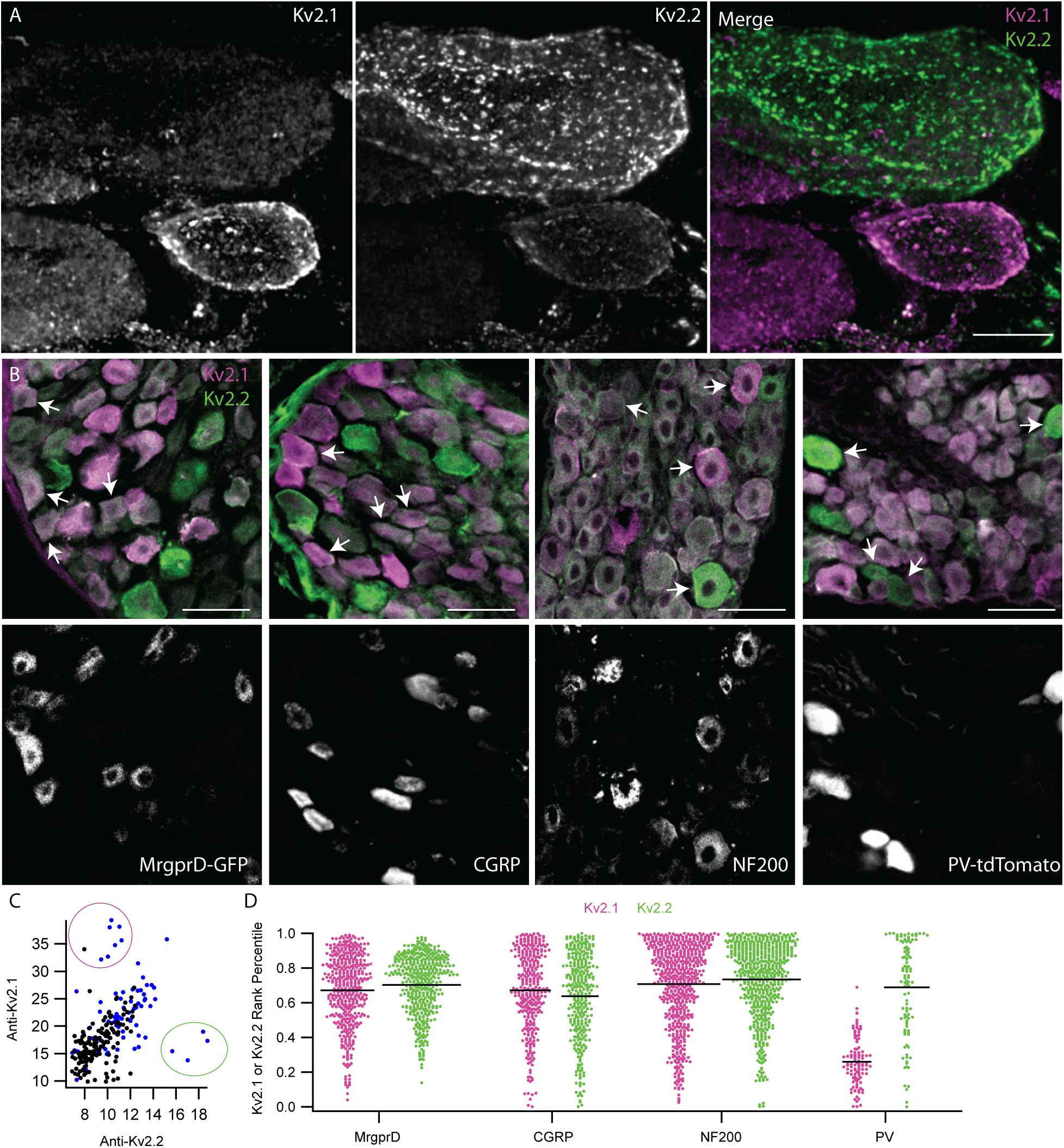
Anti-Kv2 immunofluorescence intensities are non-uniform across DRG neuron subtypes. ***A***, Exemplar z-projection of enrichment of anti-Kv2.1 (magenta) or anti-Kv2.2 (green) immunofluorescence in neighboring neurons. DRG section is from a 10 week old male mouse. Scale bar is 10 μm. ***B***, Top images show anti-Kv2.1 (magenta) and anti-Kv2.2 (green) immunofluorescence in DRG sections where subpopulation specific markers were used to identify, from left to right, non-peptidergic nociceptors, peptidergic nociceptors, myelinated neurons and proprioceptors. Fluorescence from specific markers is shown in bottom panels. Arrows indicate four exemplar neurons that have clear positivity for each subpopulation identified by fluorescence in lower panels. CGRP and NF200 subpopulations were identified using anti-CGRP and anti-NF200 antibodies while MrgprD-GFP and PV-tdTomato subpopulations were from transgenic mouse lines. Scale bars are 50 µm. ***C***, Scatter plot of anti-Kv2.1 and anti-Kv2.2 immunofluorescence of individual neuron profiles. Each point represents one profile. Magenta circle highlights the subpopulation of profiles that have high anti-Kv2.1 but low anti-Kv2.2 immunofluorescence while the green circle highlights the subpopulation of profiles that have high anti-Kv2.2 but low anti-Kv2.1 immunofluorescence. Blue points represent myelinated DRG neuron profiles identified by NF200 immunofluorescence. ***D***, Ranked anti-Kv2.1 immunofluorescence (magenta points) or ranked anti-Kv2.2 immunofluorescence (green points) of individual profiles from subpopulations shown in B. Only profiles that were positive for each marker are shown. Each point represents one profile. MrgprD population N = 4 mice, CGRP population N = 3 mice, NF200 population N = 3 mice and PV population N = 2 mice. Detailed information on each mouse used can be found in table 1.

### Kv2 channel cellular expression and subcellular localization in human DRG neurons is similar to mice

Kv2 mRNA transcripts are present in human DRG neurons (Tavares-Ferreira et al., 2022; Wangzhou et al., 2020), and we assessed whether anti-Kv2 immunofluorescence is associated with human DRG neurons. As human Kv2.1 and Kv2.2 knockout controls are not available, we compared fluorescence in DRG sections with anti-Kv2 primary antibodies to those lacking primary antibodies. Control samples lacking primary antibodies revealed strong autofluorescence at all excitation wavelengths (Supplemental Figure 11 A and B) consistent with other studies of human DRG neurons which have attributed this autofluorescence to lipofuscin (Shiers et al., 2021). We found that most neurons have anti-Kv2.1 and anti-Kv2.2 immunofluorescence brighter than controls lacking primary antibodies (Supplemental Figure 11 D and E) suggesting that Kv2 protein is present in human DRG neurons. As a control, we used the same methods with anti-Nav1.7 and anti-Nav1.8 antibodies and found anti-Nav1.7 immunofluorescence in nearly all human DRG neurons while anti-Nav1.8 was in a smaller fraction of DRG neurons, consistent with previous reports of Nav1.7 protein and Nav1.8 transcript in human DRG (Shiers, Klein, & Price, 2020) (Supplemental Figure 11 A arrow heads, B, F and G). Similar to DRG neurons in mice, we observed subpopulations of neurons that had bright anti-Kv2.1 immunofluorescence but dim anti-Kv2.2 immunofluorescence and conversely neurons that had bright anti-Kv2.2 immunofluorescence but dim anti-Kv2.1 immunofluorescence (Figure 10 A and B). These observations were consistent in DRG from 3 human donors (Supplemental Figure 11 A and H and Supplemental Figure 12). We were unable to identify if the population of human DRG neurons with low anti-Kv2.1 and high anti-Kv2.2 are proprioceptors as we observed in mice; attempts to label parvalbumin in human neurons were unsuccessful.

**Figure 10.**
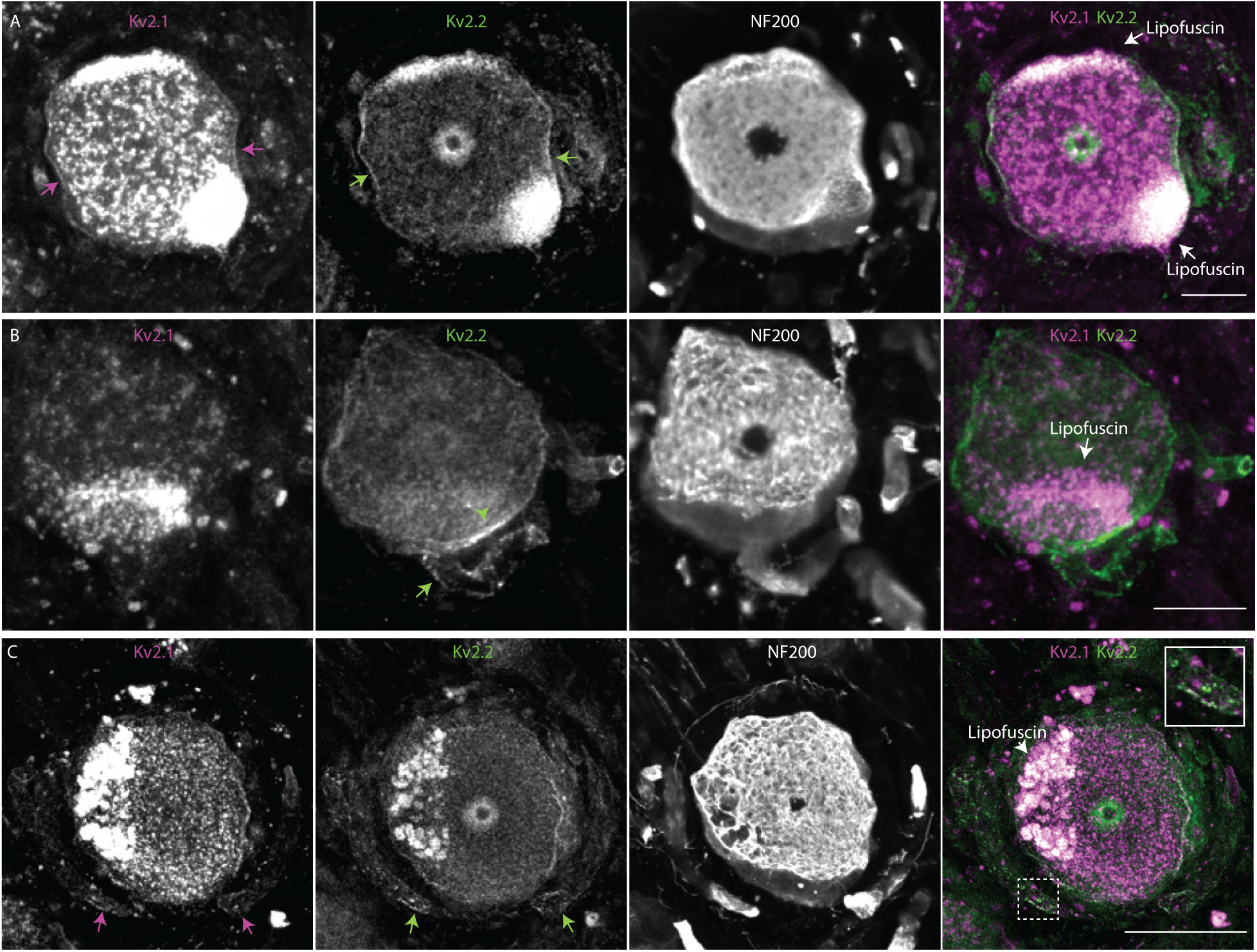
Kv2 channel expression and localization in human DRG neurons is similar to mice. ***A***, Immunofluorescence from human DRG neurons labeled with anti-Kv2.1 and anti-Kv2.2 antibodies. Autofluorescence attributed to lipofuscin is labeled in right panel while apparent Kv2.1 and Kv2.2 protein are labeled in left and middle panel respectively. Scale bar is 50 μm. ***B***, Z-projection of anti-Kv2.2 (left) and anti-NF200 immunofluorescence (middle) of human DRG neuron somata. Green arrow head indicates asymmetric distribution of Kv2.2 clusters on neuron soma, green arrow indicates the apparent stem axon. Scale bar is 20 μm. ***C***, Z-projection of anti-Kv2.1 (upper left), anti-Kv2.2 (upper right) and anti-NF200 (lower left) immunofluorescence of a human DRG neuron. Magenta and green arrows indicate Kv2.1 and Kv2.2 respectively on the apparent stem axon. Inset shows expansion of dotted line boxes which highlights Kv2.1 and Kv2.2 clusters on the apparent stem axon. Autofluorescence attributed to lipofuscin is labeled in lower right panel. Scale bar is 50 μm. All images are from donor #1. Detailed information on each donor can be found in the *Human Tissue Collection* section of the methods.

We assessed whether Kv2 protein subcellular localization in humans is similar to mice. We identified neurons in all three human DRG samples where anti-Kv2.1 and anti-Kv2.2 immunofluorescence was enriched at the outer edge of DRG neuron somata (Figure 10 A arrow and Supplemental Figure 13). In some neurons from all three humans, Kv2 clusters exhibited asymmetric distribution on the neuronal soma (Figure 10 B green arrow head, Supplemental Figure 13 A-E arrows). In one human we identified a neuron with Kv2.2 immunofluorescence that was asymmetrically distributed near the stem axon consistent with observations in mice (Figure 10 B green arrow). We identified Kv2.2 immunofluorescence on the stem axon of DRG neurons in all three humans using anti-NF200 or anti-Nav1.7 as markers of the stem axon (Figure 10 B-C and Supplemental Figure 14 arrows). In two out of three humans we also identified Kv2.1 immunofluorescence on the stem axon of human DRG neurons (Figure 10 C inset and Supplemental Figure 14 B and D inset). To determine if Kv2.2 channels are present in human DRG neuron axons, we labeled human DRG sections with the same anti-Kv2.2, anti-NF200, anti-CASPR and anti-Kv1.2 antibodies that were used in mice. We found anti-Kv2.2 immunofluorescence associated with anti-NF200, anti-CASPR and anti-Kv1.2 immunofluorescence in human DRG neuron axons (Supplemental Figure 15) suggesting that Kv2.2 channels could contribute to electrical propagation or other physiological functions in axons of human DRG neurons.

## DISSCUSSION

Our results reveal overlapping yet distinct localization patterns of Kv2.1 and Kv2.2 in DRG neurons. Here, we discuss the unique features of Kv2 localization in the DRG and how these could influence neuronal physiology and somatosensation.

We identify differences between anti-Kv2.1 and anti-Kv2.2 immunofluorescence in DRG neurons. Kv2.2 mRNA is abundant in all DRG neuron subtypes identified, while Kv2.1 mRNA is low in proprioceptors and abundant in other subtypes (Usoskin et al., 2015; Zheng et al., 2019). Consistent with these findings we conservatively estimate that Kv2.1 and Kv2.2 protein is in at least 90% of mouse DRG neurons, though Kv2.1 immunofluorescence is anomalously low in proprioceptors. Detection of Kv2.2 but not Kv2.1 in DRG neuron peripheral and central axons suggests that Kv2.2 channels may be important for electrical transmission in these axons. While detectable Kv2.2 remained constant in DRG neurons of older mice, detectable Kv2.1 decreased, possibly contributing to the altered repolarization noted in DRG neurons from aged mice (Scott, Leu, & Cinader, 1988). The differences between Kv2.1 and Kv2.2 in somatosensory neurons are consistent with distinct roles for these Kv2 paralogs in somatosensory neuron physiology, as seen in brain and pancreatic islets (Bishop et al., 2015; Johnston et al., 2008; Li et al., 2013).

The expression and subcellular localization of Kv2 channels in DRG neurons are distinct from central neurons. Kv2.1 is present in most brain neurons (Bishop et al., 2018; Bishop et al., 2015; Hwang, Fotuhi, Bredt, Cunningham, & Snyder, 1993) while Kv2.2 is less broadly expressed (Bishop et al., 2015; Hermanstyne et al., 2010; Hwang, Glatt, Bredt, Yellen, & Snyder, 1992; Johnston et al., 2008; Kihira et al., 2010). In contrast to this, we find Kv2.2 expressed in at least 90% of mouse DRG neurons. We also identify that Kv2.2 is enriched in the DRG relative to neurons in the spinal cord, suggesting that abundant Kv2.2 protein expression is a distinct feature of DRG neuron physiology. In central neurons, Kv2 channel clusters are typically evenly distributed across neuron somata (Bishop et al., 2015; Kihira et al., 2010; Muennich & Fyffe, 2004) and asymmetric subcellular localization of Kv2 clusters on neuron somata has to our knowledge not been described. We find that Kv2 channels are enriched at the plasma membrane in a subcellular region near the stem axon of some DRG neurons. Differences in Kv2 localization between central and somatosensory neurons could arise from differences in the extracellular environment surrounding DRG neuron somata. In central neurons, Kv2 channel clusters are reported at cholinergic C-terminal synaptic sites, S-type synapses, and apposed to astrocytic end feet (Du et al., 1998; Muennich & Fyffe, 2004). However, synapses have not been reported in the DRG and DRG neurons are surrounded by satellite glia instead of astrocytes (Matsuda et al., 2005). Further studies could investigate if Kv2 clusters are localized to extracellular structures in the DRG. Kv2.2 has been identified in juxtaparanodes but not paranodes of axons in the osseous spiral lamina of the cochlea (Kim & Rutherford, 2016) and previous work has also identified that Kv2.1 channels are necessary for enabling ER Ca2+ uptake during electrical activity in both the soma and axon (Panzera et al., 2022). To our knowledge, Kv2.2 immunofluorescence has not been previously reported in distal axons in the brain, spinal cord or DRG. Our results indicate that Kv2.2 channels are present in myelinated and unmyelinated axons of DRG neurons. The mechanism of Kv2.2 protein trafficking to the axon is unknown. Kv2.1 channels are trafficked to the axon initial segment through a distinct secretory pathway (Jensen et al., 2017) allowing neurons to regulate precise compartment specific localization of Kv2.1 protein. A similar mechanism for Kv2.2 localization in DRG axons could allow neurons to independently control Kv2.2 protein expression in the neuron soma and axon. The subcellular distribution of Kv2.1 and Kv2.2 subunits in DRG neurons suggests they could each distinctly modulate electrical signals. We note that our observations could not distinguish whether immunofluorescence in DRG axons could represent intracellular trafficking vesicles of Kv2.2 proteins, or whether axonal Kv2.2 is on the membrane surface where it could modulate electrical signals.

Kv2 channels in DRG neurons may play important roles in somatosensation and nociception. Kv2 conductances can modulate action potential repolarization, afterhyperpolarization and repetitive firing (Liu & Bean, 2014; Zheng et al., 2019). Kv2 conductances modulate electrical transmission in rat DRG neurons (Tsantoulas et al., 2014). Knockdown of Kv2.1 increases hypersensitivity to painful stimuli and regulation of Kv2.1 protein expression through the epigenetic factor Cdyl modulates neuronal excitability and nociception (Sun et al., 2022). Consistent with these studies, our results show that Kv2 channels are localized to DRG neuron axons, where they could potentially influence electrical transmission.

Regulatory pore-forming subunits of the Kv5, Kv6, Kv8, and Kv9 “silent subunit” families obligately assemble into heteromeric channels with Kv2 subunits (Bocksteins, 2016) and have been implicated in nociception. Women with decreased sensitivity to labor pain were identified to have a mutation in the Kv6.4 silent subunit that disrupts Kv2.1/Kv6.4 heteromer expression and is proposed to modulate nociceptor signaling (Lee et al., 2020). Downregulation of the Kv9.1 silent subunit after axotomy evokes hyperexcitability of DRG neurons (Tsantoulas et al., 2012). While both Kv2 channels are broadly expressed in DRG neurons, transcript levels of silent subunits in mice and humans show much more neuron subtype specificity (Tavares-Ferreira et al., 2022; Zheng et al., 2019), raising the possibility of molecularly distinct Kv2-containing channels in somatosensory neuron subtypes.

Previous studies have identified that voltage gated ion channel expression can be substantially different between mice and human DRG, notably, in the expression of Nav1.7 (Shiers et al., 2020). However, we find expression of Kv2 channels to be similar in mouse and human DRG. An overall similarity is supported by the presence of detectable Kv2 channels in most neurons, consistent with transcriptomics data from both mice and humans (Tavares-Ferreira et al., 2022; Zheng et al., 2019). The subcellular localization of Kv2 channels also appears similar between mice and humans. These similarities include enrichment of Kv2 channels at the outer edge of neurons, Kv2 clusters on the neuron soma as well as presence of Kv2.1 and Kv2.2 on DRG neuron axons. These similarities suggest that the functional roles of Kv2 channels could be similar in both mouse and human DRG neurons.

Overall, we find attributes of Kv2 channel localization in somatosensory neurons that could point to their functional roles in sensory information processing. The broad expression of Kv2.2 among DRG neuron types and expansive axonal localization of Kv2.2 suggest a key role for this channel in the somatosensory nervous system. The specialized divergence of Kv2.1 and Kv2.2 suggest the functional roles of Kv2 subtypes in DRG neurons are overlapping yet distinct.

## Data availability

The data that support the findings of this study are available upon request from the authors. Code used to analyze this data in ImageJ and R is available at https://github.com/SackLab/DRG-Image-Processing

## Acknowledgments

We thank James S. Trimmer (University of California, Davis) for generous provision of antibodies, numerous discussions, and feedback on the manuscript. We thank Michael Ferns (University of California, Davis) for feedback on the manuscript. We thank the human tissue donors and their families for their generous donations to science. We thank Jiwon Yi, Zack Bertels, Maria Payne, and Jon Lemen for performing hDRG extractions. GxTX was synthesized at the Molecular Foundry of the U.S. Department of Energy under contract DE-AC02-05CH11231. This research was supported by U.S. National Institutes of Health grant R01NS096317 to J.T. Sack and the University of California, Davis.

## Author contributions

R. G. Stewart: Conceptualization, formal analysis, investigation, methodology, visualization, writing-original draft, writing-reviewing and editing. M. Camacena: investigation. B. A. Copits investigation, writing-reviewing and editing. J.T. Sack: Conceptualization, formal analysis, funding acquisition, investigation, methodology, project administration, supervision, visualization, writing- original draft, writing-reviewing and editing.

## SUPPLEMENTAL FIGURES

**Supplemental Figure 1.**
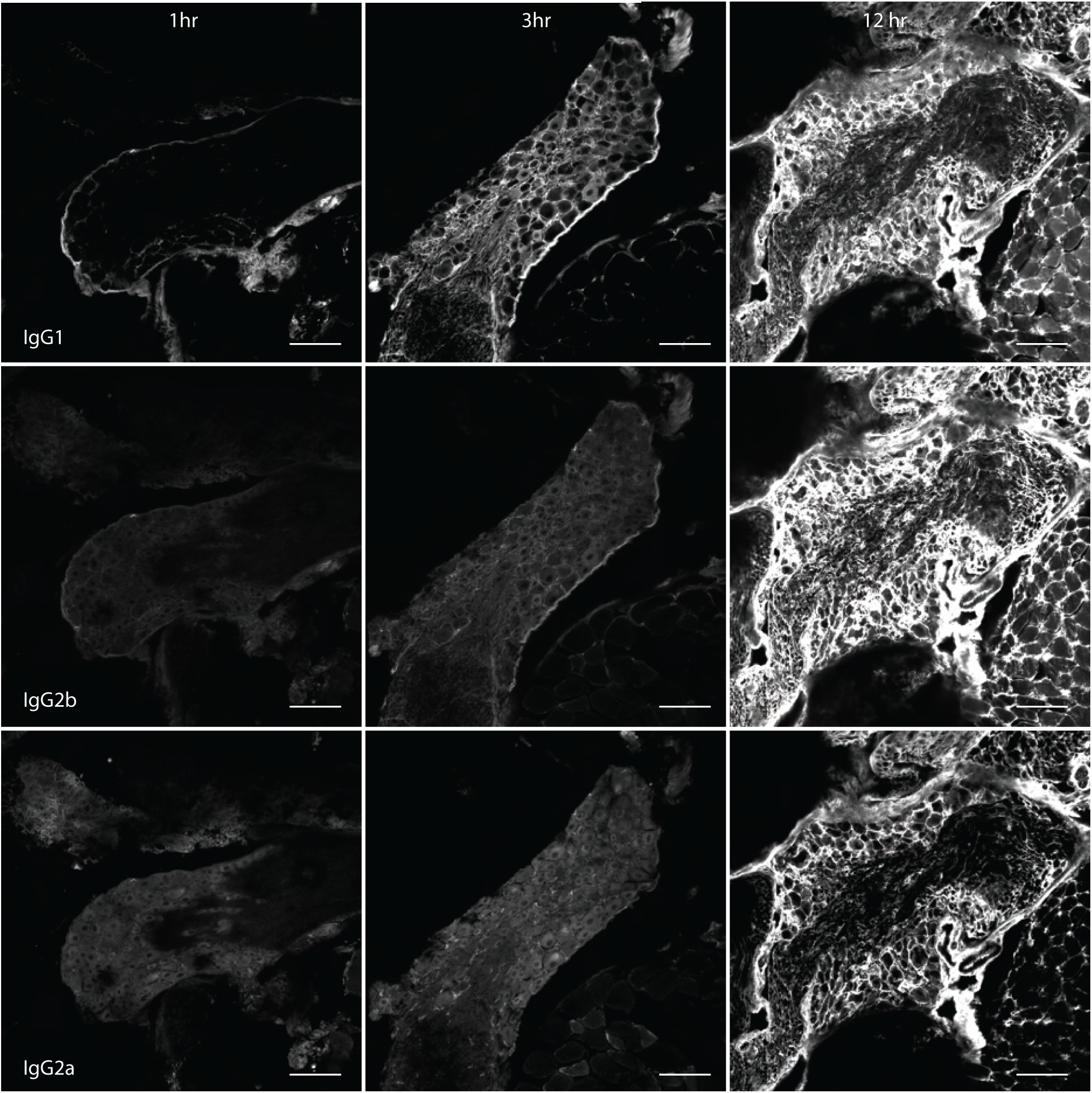
Off-target mouse anti-IgG1, IgG2b, and IgG2a immunofluorescence increases with fixation time. Images of DRG sections labeled with fluorescently tagged antibodies which target IgG1 (top), IgG2b (middle) and IgG2a (bottom) after DRG were fixed in ice cold 4% PFA for 1, 3 and 12 hours. DRG sections are from the same mouse. Scale bars are 100 μm.

**Supplemental Figure 2.**
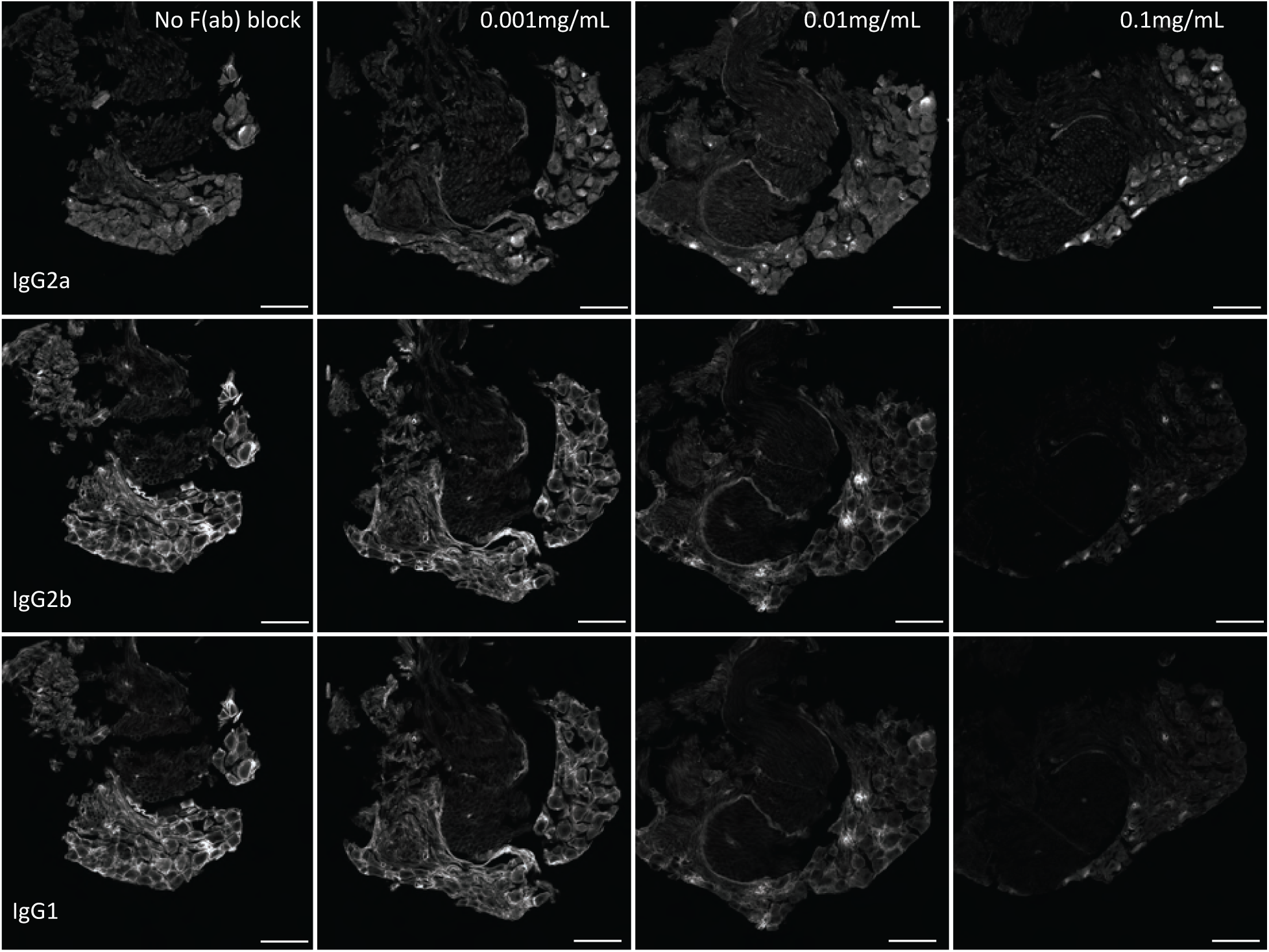
Pre-incubation of mouse DRG sections in IgG H+L Fab fragments reduces off target secondary antibody labeling. Representative images of DRG sections from the same DRG treated with increasing concentrations (left to right) of IgG H+L Fab fragment and the same concentration of secondary antibody used in experiments throughout this study. Images were taken with identical imaging settings and are set to the same brightness and contrast. Scale bars are 100 μm.

**Supplemental Figure 3.**
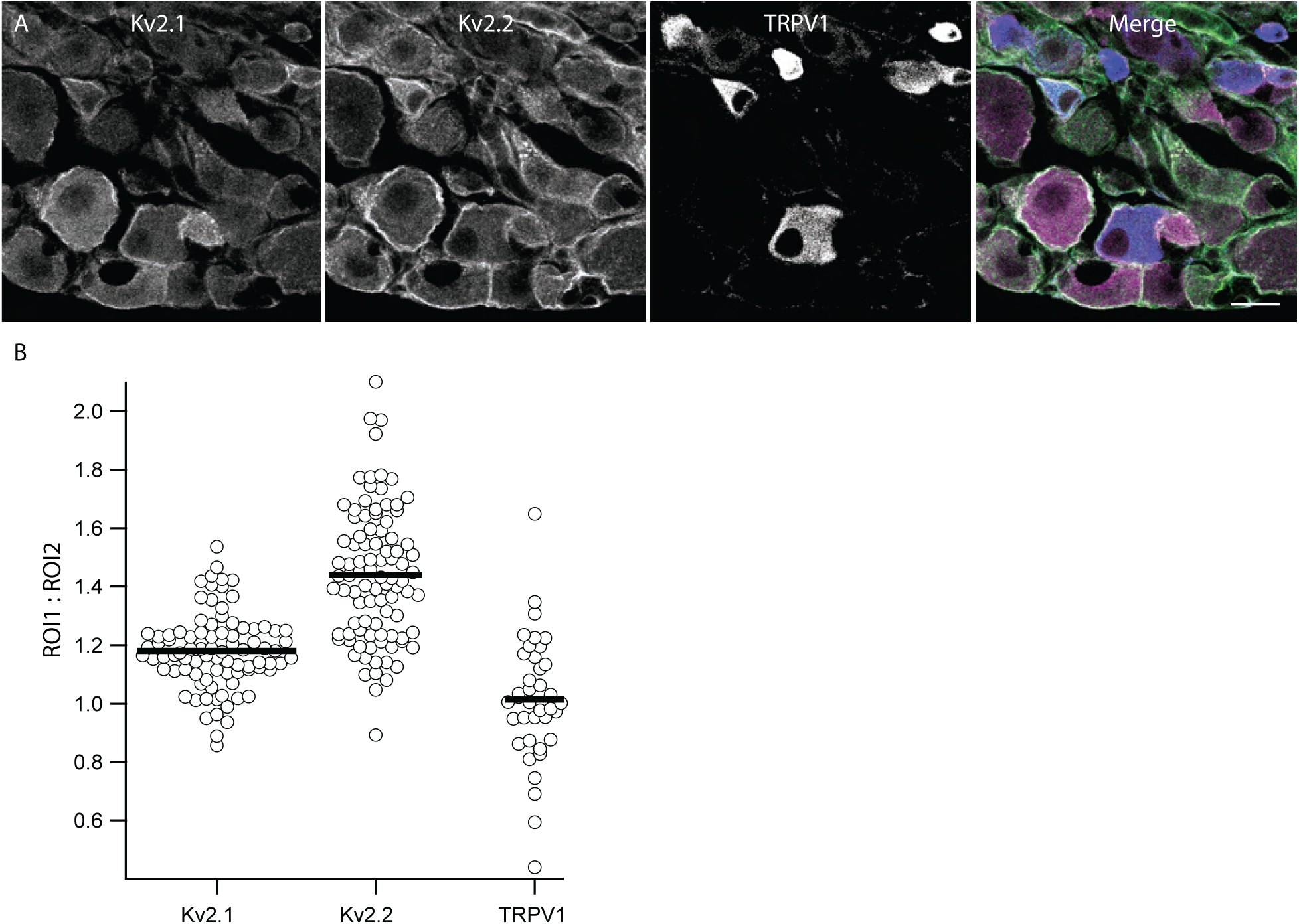
Kv2.1 and Kv2.2 protein are enriched at the outer edge of DRG neuron somas relative to TRPV1. ***A***, Anti-Kv2.1, anti-Kv2.2 and anti-TRPV1 immunofluorescence from lumbar DRG neurons. Prominent cytoplasmic anti-TRPV1 immunofluorescence was observed in a subset of small diameter neurons. In merge image anti-Kv2.1, anti-Kv2.2 and anti-TRPV1 immunofluorescence are magenta, green and blue respectively. Scale bar is 20 μm. ***B***, Ratio of average anti-Kv2.1, anti-Kv2.2 or anti-TRPV1 immunofluorescence from outer and inner ROIs for individual neurons.

**Supplemental Figure 4.**
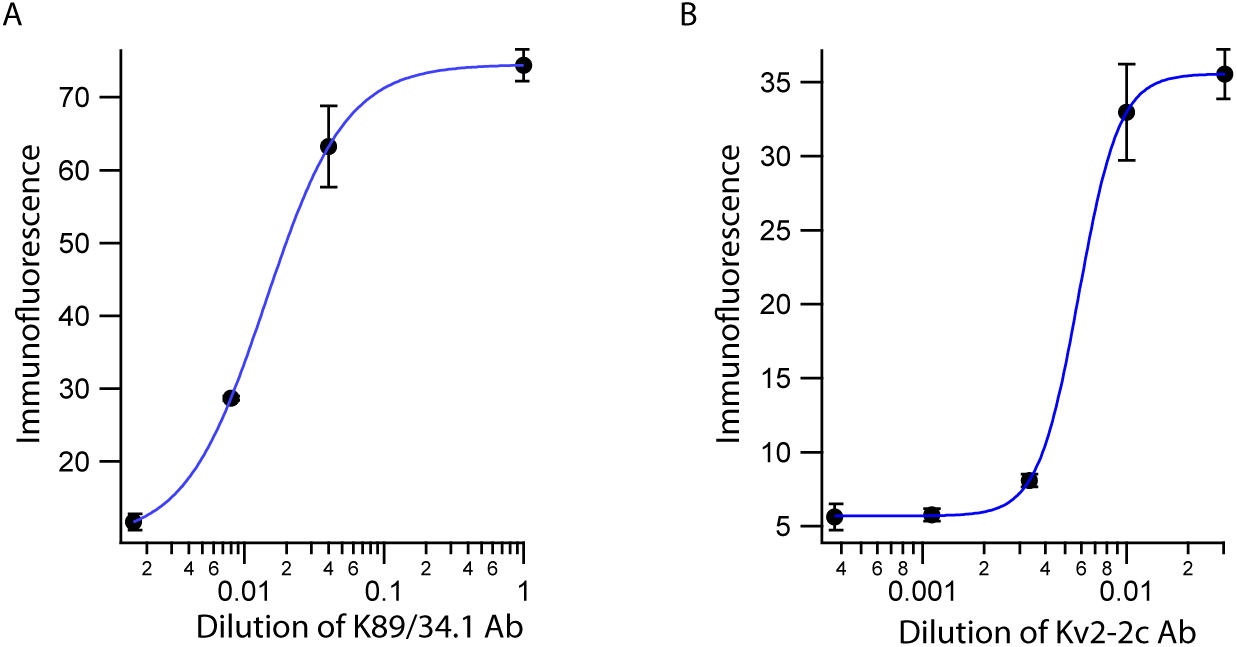
Kv2 antibodies used in knockout experiments are at saturating concentrations. ***A***, Concentration response of immunofluorescence from sections labeled with anti-Kv2.1 antibody used in Figure 2. Blue line is a Hill fit of the data. 1:1 = 2 sections, 1:25 = 3 sections, 1:125 = 2 sections, 1:625 = 3 sections ***B***, Concentration response of immunofluorescence from sections labeled with anti-Kv2.2 antibody used in Figure 3. Blue line is a Hill fit of the data. 1:33 = 2 sections, 1:100 = 10 sections, 1:300 = 4 sections, 1:900 = 4 sections, 1:2700 = 3 sections

**Supplemental Figure 5.**
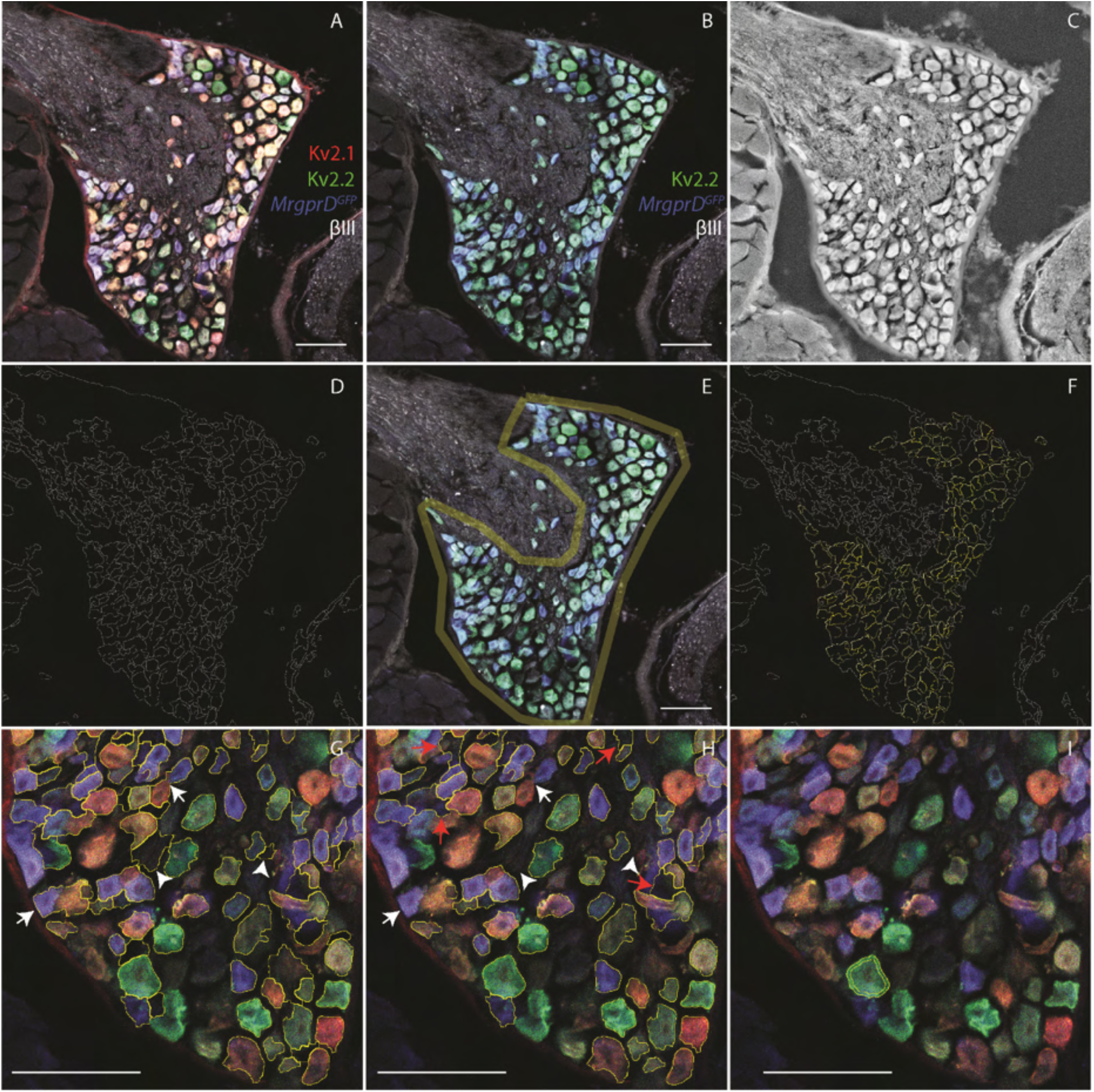
A method to sample neurons in DRG imaging data using watershed segmentation identifies the outer region of neurons. ***A***, Image of anti-Kv2.1, anti-Kv2.2, anti-βIII tubulin immunofluorescence and MrgprD-GFP fluorescence. ***B***, Same image as A with anti-Kv2.1 immunofluorescence channel removed as this is an example of processing data for Kv2.1 KO analysis. ***C***, Grayscale image of average fluorescence from all three channels shown in B. Gaussian and median filters were applied to image to improve watershed segmentation. ***D***, Watershed segmentation of image in C using the MorphoLibJ Morphological Segmentation Plugin in Fiji. ***E***, Example of manually drawn boundary that encompasses the neuron somas in the DRG so that only these watershed lines are selected. ***F***, Selected ROIs from watershed segmentation (yellow). ROIs were excluded based on roundness and size using the Analyze Particles tool in Fiji. ***G***, ROIs in F overlaid on DRG image showing that some ROIs are selecting regions that do not contain neurons (arrow heads) or are selecting multiple neurons (arrows). ***H***, ROIs after processing using an in-house R script which removes ROIs that do not contain neurons (arrow heads) and ROIs that contain two neurons (arrows). This script did not remove all ROIs that do not contain neurons (red arrows). Each experiment performed was done alongside controls where the primary antibodies were omitted and fluorescence from these control sections was used by the in-house R script to identify and remove ROIs that do not contain neurons. ***I***, Example of automatically generated annulus that encompasses the outer edge of the soma. Scale bars are 100 µm.

**Supplemental Figure 6.**
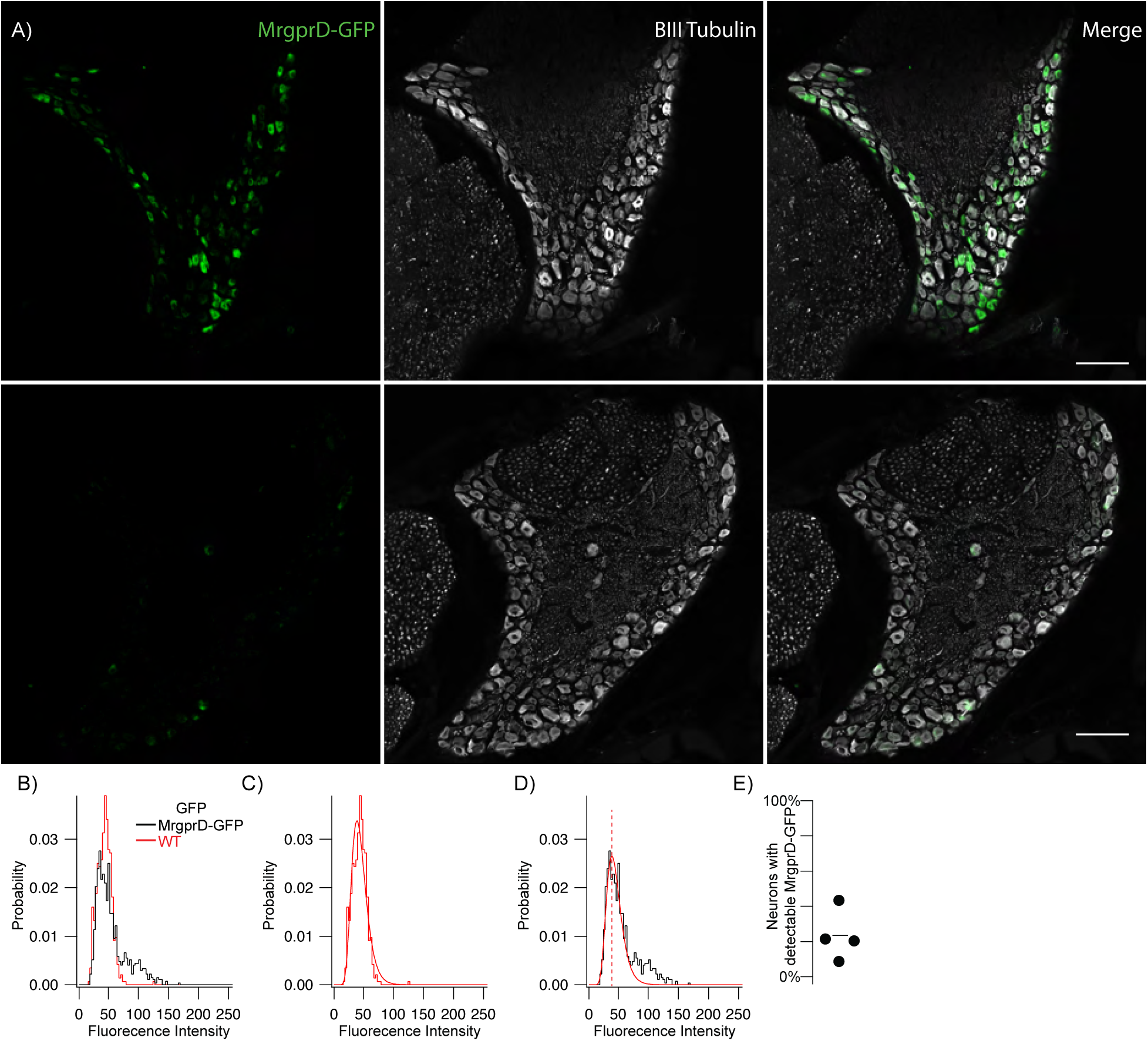
Method used to estimate percent of neurons expressing Kv2.1 and Kv2.2 reliably predicts the percentage of neurons that express GFP in MrgprD-GFP mice. ***A***, MrgprD-GFP (top) and WT (bottom) DRG sections immunolabeled for BIII tubulin (white). Images were taken with identical imaging settings and are set to the same brightness and contrast. Scale bars are 100 µm. ***B***, Distribution of fluorescence intensity from MrgprD-GFP (black) and WT (red) neurons. Data represents the fluorescence intensity of 905 MrgprD-GFP neurons from 9 DRG sections from 1 mouse or 477 WT neurons from 5 DRG sections from 1 mouse. DRG sections were taken from 7 week old female mice and are from the 1^st^ lumbar DRG. ***C***, WT data shown in B fit with a log normal distribution (red fit). ***D***, MrgprD-GFP data shown in B fit with the WT distribution (red fit) where width and mean were constrained to the WT distribution and amplitude was unconstrained (equation 1). Only MrpgrD-GFP data to the left of the mean intensity of WT neurons (red dotted line) was used for the fit. ***E***, Percent of neurons with detectable GFP protein of 4 mice (3 females 1 male). All DRG sections were taken from the 1^st^ lumbar DRG.

**Supplemental Figure 7.**
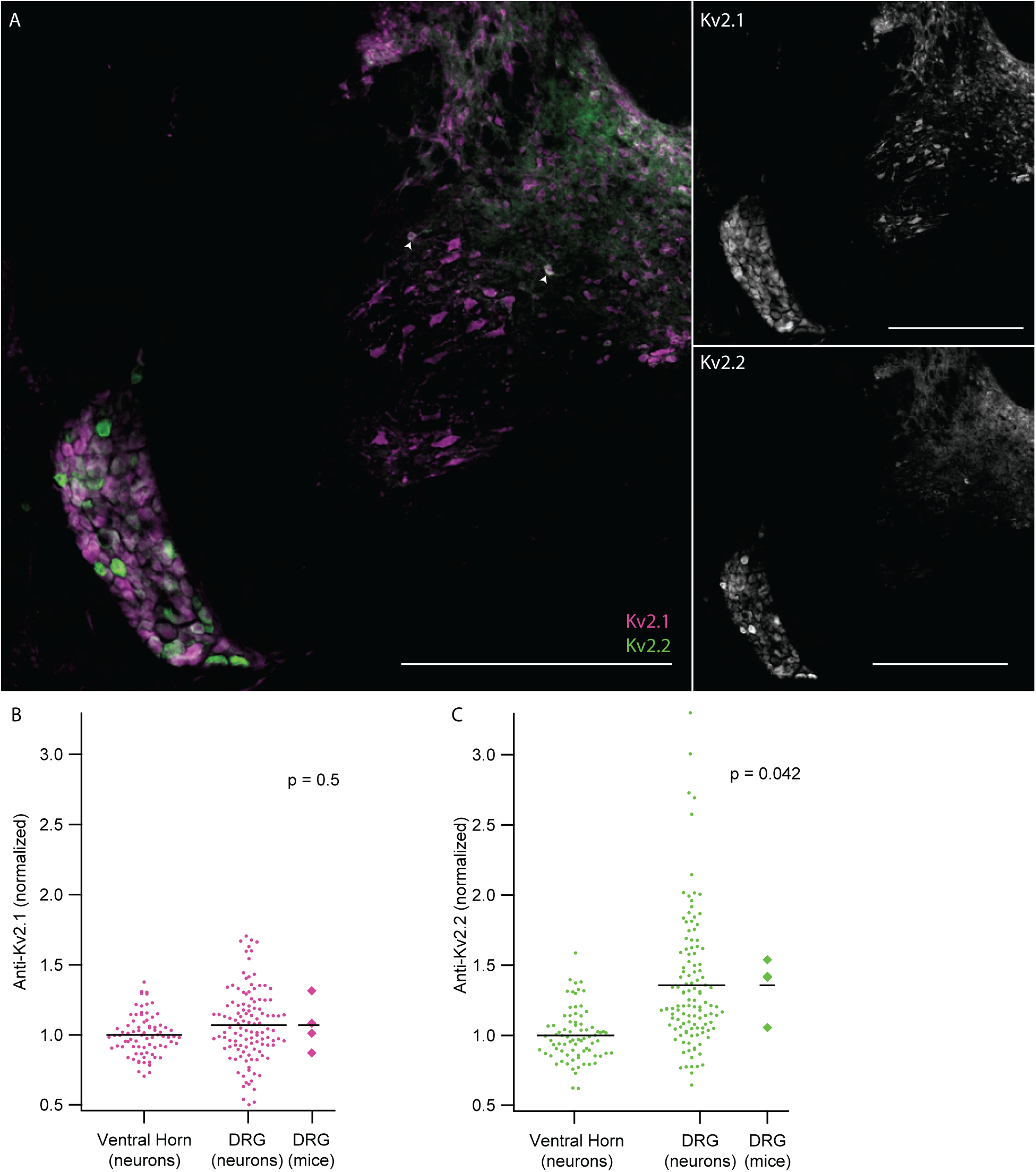
DRG neurons have enriched Kv2.2 protein compared to neurons in the ventral horn. ***A***, Anti-Kv2.1 (magenta) and anti-Kv2.2 (green) immunofluorescence in a spinal cord section from the 13^th^ thoracic vertebra (left). Anti-Kv2.1 immunofluorescence (right top) and anti-Kv2.2 immunofluorescence (right bottom). Arrow heads show neurons in the spinal cord with anti-Kv2.2 immunofluorescence. Scale bars are 500 μm. ***B***, Anti-Kv2.1 immunofluorescence from individual neurons (circles) in the DRG and ventral horn normalized to the average fluorescence intensity of neurons in the ventral horn. Diamonds to the right of data represent the average intensity in the DRG of individual mice. Significant differences from 1 were calculated for individual mice using Students t-test. N = 4 mice n = 116 in DRG and n = 77 in ventral horn. ***C***, Identical analysis shown in B with anti-Kv2.2 immunofluorescence.

**Supplemental Figure 8.**
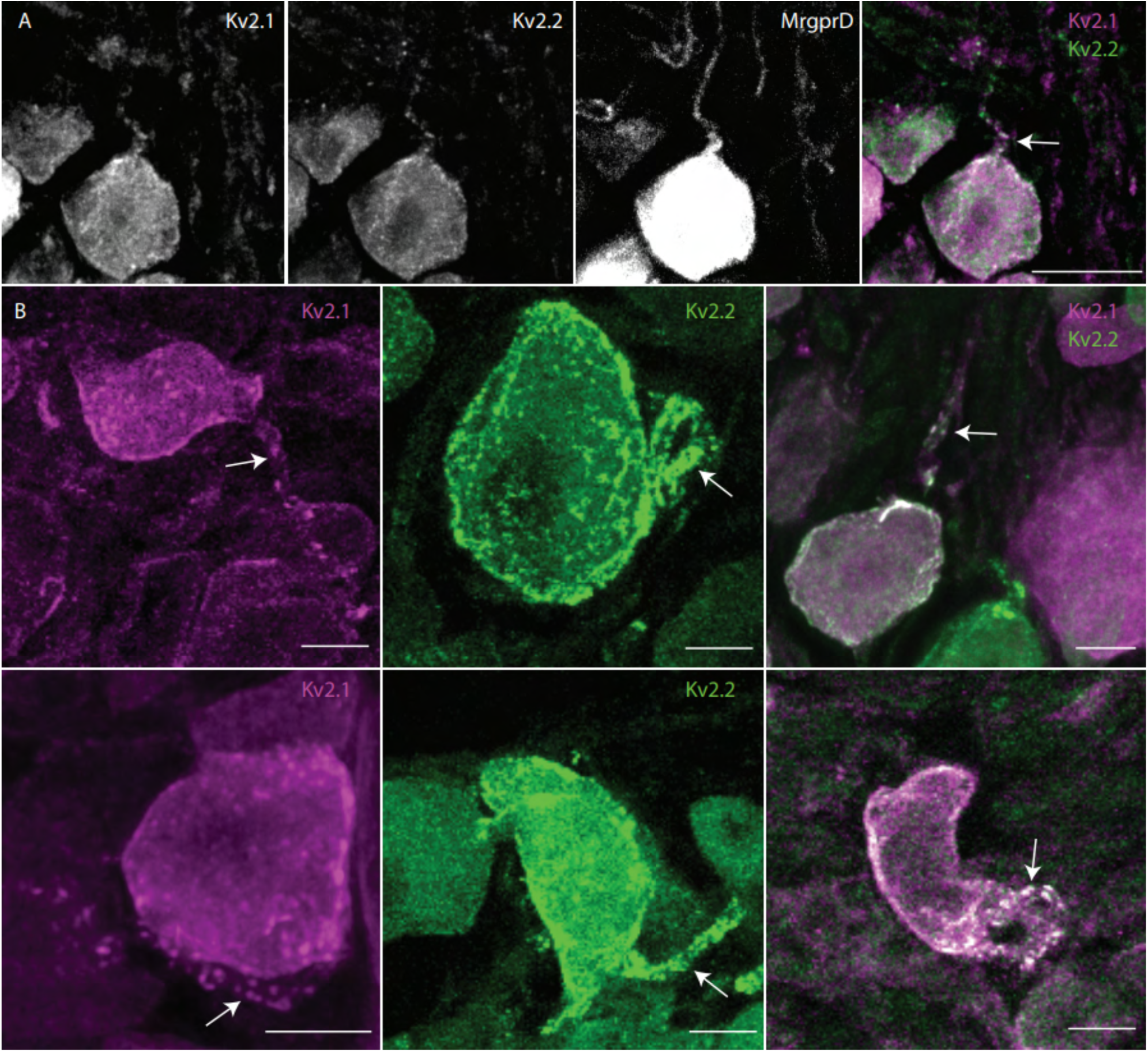
Kv2 channels are expressed on the stem axon of mouse DRG neurons. ***A***, Z-projection with anti-Kv2.1 and anti-Kv2.2 immunofluorescence on the stem axon of a neuron in the DRG of a MrgprD-GFP mouse. ***B***, Gallery of z-projected images of DRG neurons with anti-Kv2.1 and/or anti-Kv2.2 immunofluorescence on stem axons. Arrows indicate stem axons. Scale bars are 10 µm

**Supplemental Figure 9.**
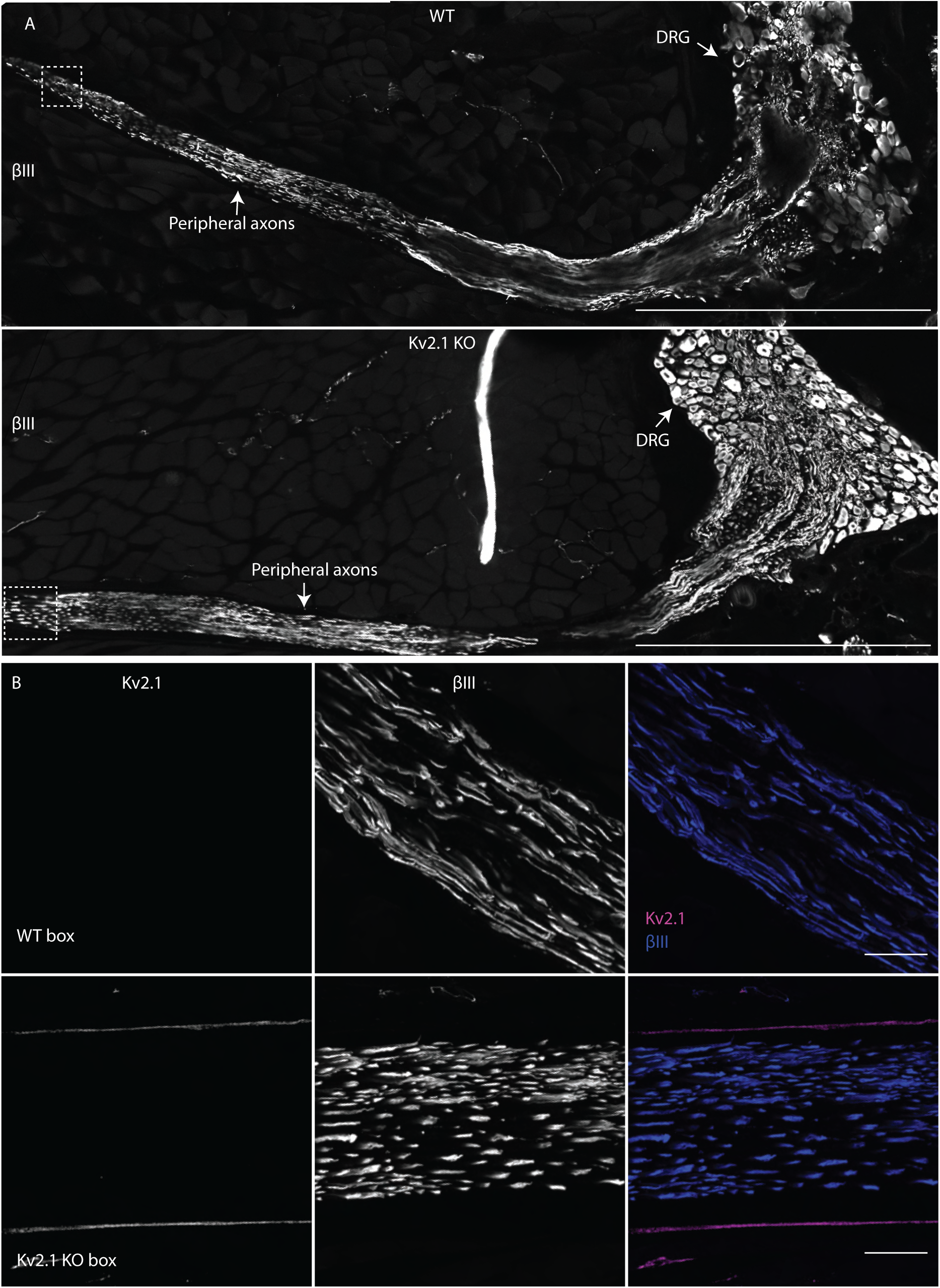
Kv2.1 channels were not detected in peripheral axons of DRG neurons. ***A***, WT (top) and Kv2.1 KO (bottom) sections containing the DRG and peripheral axons from the 12^th^ thoracic DRG in age and sex matched 7 week old mice immunolabeled for βIII tubulin (white). Scale bar is 500 µm. ***B***, High magnification z-projection of anti-Kv2.1 and anti-βIII immunofluorescence from box in A of WT and Kv2.1 KO mice. Scale bars are 20 µm.

**Supplemental Figure 10.**
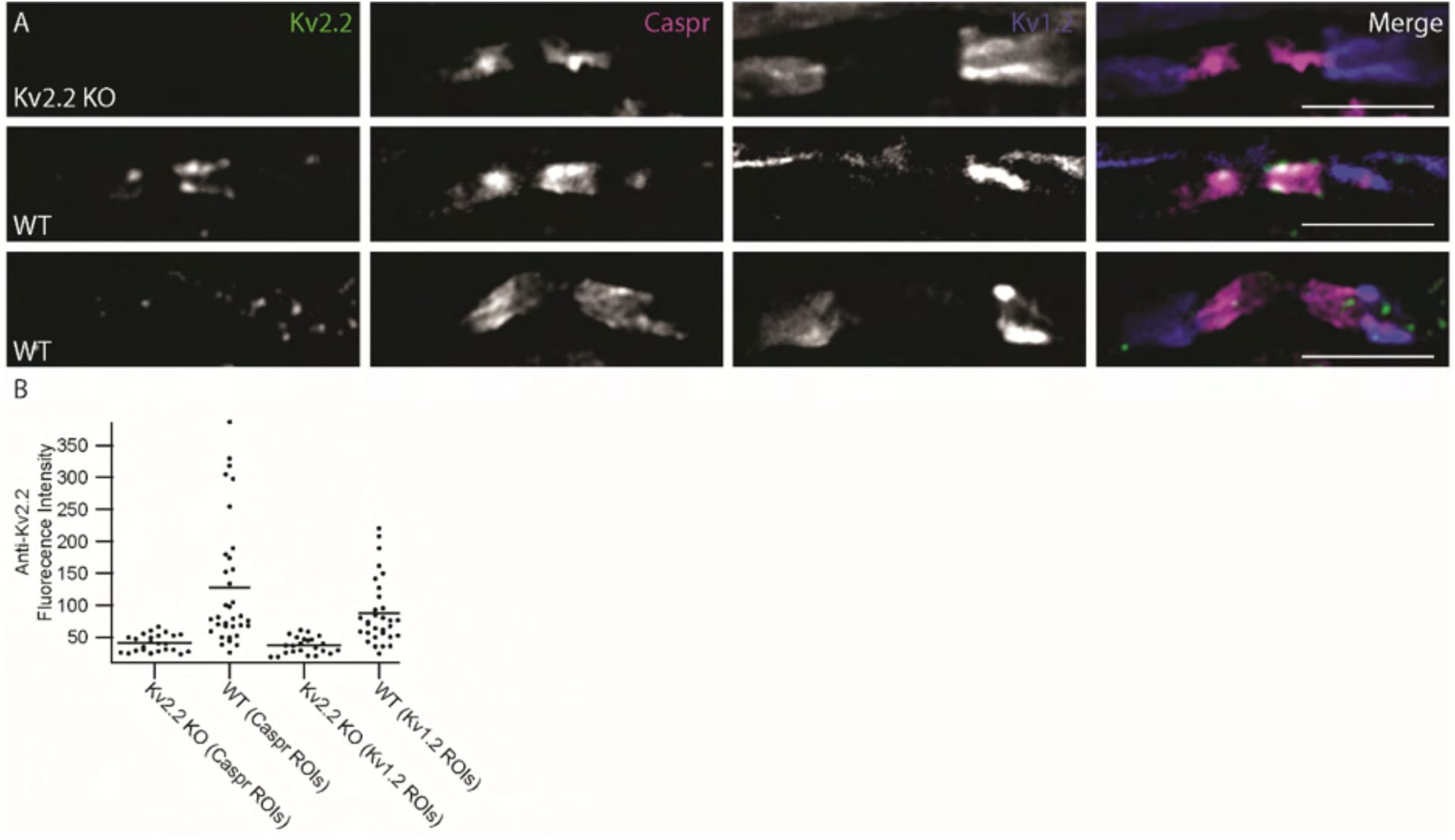
Kv2.2 is expressed in myelinated fibers of DRG neuron axons. ***A***, Kv2.2 KO (top) and WT (middle and bottom) sections containing the peripheral axons from the 12^th^ thoracic DRG in 28 week old mice immunolabeled for Kv2.2, Caspr and Kv1.2. Middle panels are an exemplar of prominent Kv2.2 immunofluorescence in CASPR labeled axons and bottom panels are an exemplar of prominent Kv2.2 clusters in the Kv1.2 labeled axons. Scale bars are 5 μm. ***B***, Analysis of anti-Kv2.2 immunofluorescence intensity in CASPR and Kv1.2 labeled regions of age and sex matched WT and Kv2.2 KO mice. Individual points represent single ROIs drawn around anti-CASPR or anti-Kv1.2 immunofluorescence.

**Supplemental Figure 11.**
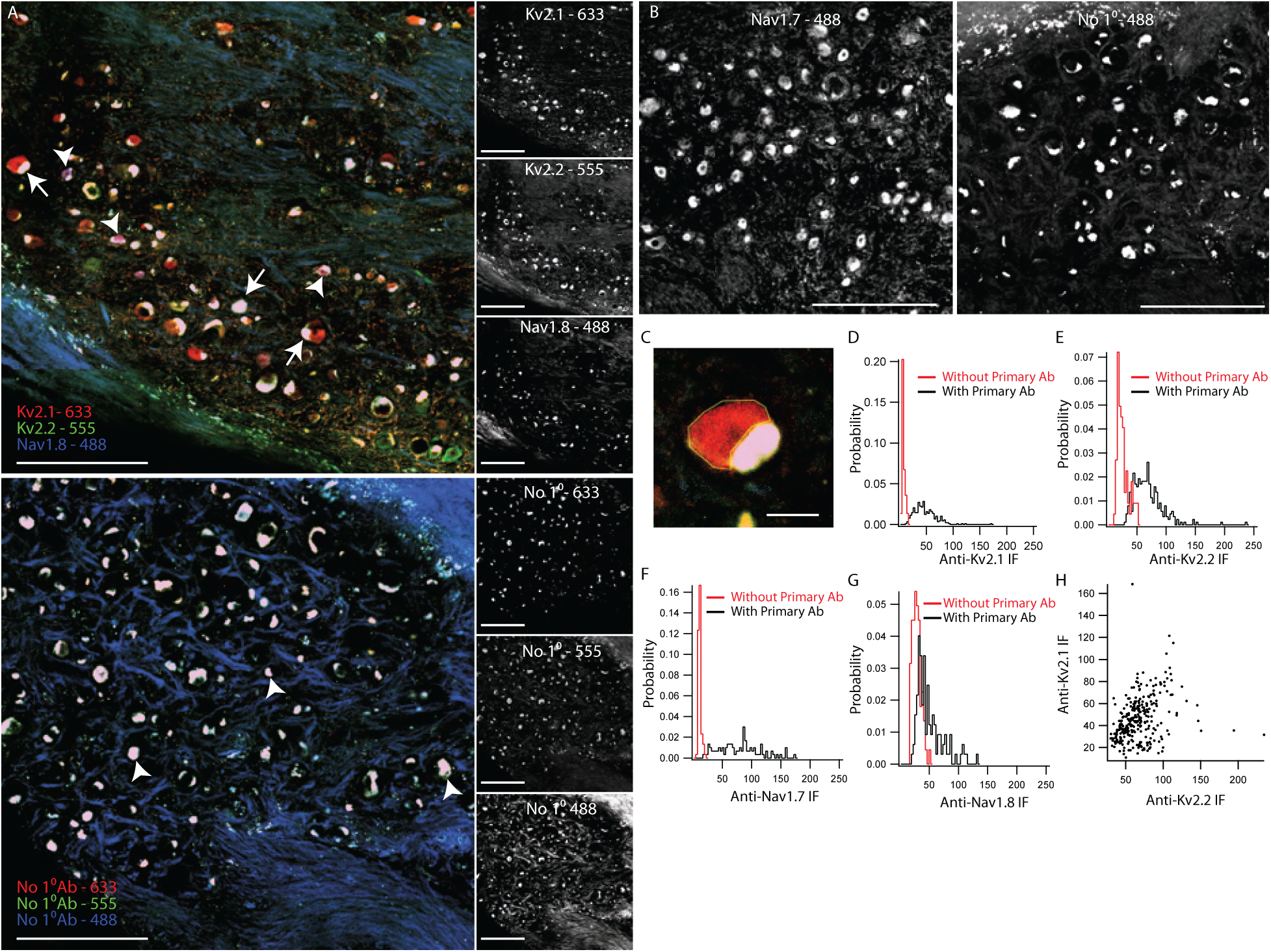
Fluorescence from human DRG neurons labeled with ion channel targeting antibodies is distinct from human DRG neurons where ion channel targeting antibodies were omitted. ***A***, Top: Immunofluorescence from human DRG section labeled with anti-Kv2.1, anti-Kv2.2 and anti-Nav1.8 antibodies. Bottom: Fluorescence from human DRG section where the primary antibodies are omitted. Arrows in top and bottom images indicate examples of autofluorescence from apparent intracellular lipofuscin. Arrow heads in top image identify anti-Nav1.8 immunofluorescence. Images on the right are fluorescence from each fluorescence channel of the top and bottom images. Number next to target protein label represents excitation wavelength. DRG sections from top and bottom images are from the same DRG. Scale bars are 500 μm. ***B***, Left: Immunofluorescence from human DRG section labeled with anti-Nav1.7 antibody. Right: Fluorescence from human DRG section where the primary antibody has been omitted. Number next to target protein label represents excitation wavelength. DRG sections in left and right images are from the same DRG. Scale bars are 500 μm. ***C***, Exemplar manually drawn ROI to analyze fluorescence intensity in human DRG neurons that omits apparent lipofuscin autofluorescence. Scale bar is 50 μm. ***D***, Distribution of fluorescence intensity of human DRG neurons labeled with an anti-Kv2.1 antibody (black) or when the anti-Kv2.1 antibody was omitted (red). Data represents the fluorescence intensity of 293 neurons labeled with anti-Kv2.1 antibody or 73 neurons where the anti-Kv2.1 antibody was omitted. ***E***, Distribution of fluorescence intensity of human DRG neurons labeled with anti-Kv2.2 antibody (black) or when the anti-Kv2.2 antibody was omitted (red). Data represents the fluorescence intensity of 293 neurons labeled with anti-Kv2.2 antibody or 73 neurons where the anti-Kv2.2 antibody was omitted. ***F***, Distribution of fluorescence intensity of human DRG neurons labeled with anti-Nav1.7 antibody (black) or when the anti-Nav1.7 antibody was omitted (red). Data represents the fluorescence intensity of 99 neurons labeled with anti-Nav1.7 antibody or 99 neurons where the anti-Nav1.7 antibody was omitted. ***G***, Distribution of fluorescence intensity of human DRG neurons labeled with anti-Nav1.8 antibody (black) or when the anti-Nav1.8 antibody was omitted (red). Data represents the fluorescence intensity of 293 neurons labeled with anti-Nav1.8 antibody or 73 neurons where the anti-Nav1.8 antibody was omitted. ***H***, Fluorescence intensity of human neurons labeled with both anti-Kv2.1 and anti-Kv2.2 antibodies. Individual points represent individual neurons. All images are from donor #2. Detailed information on each donor can be found in the *Human Tissue Collection* section of the methods.

**Supplemental Figure 12.**
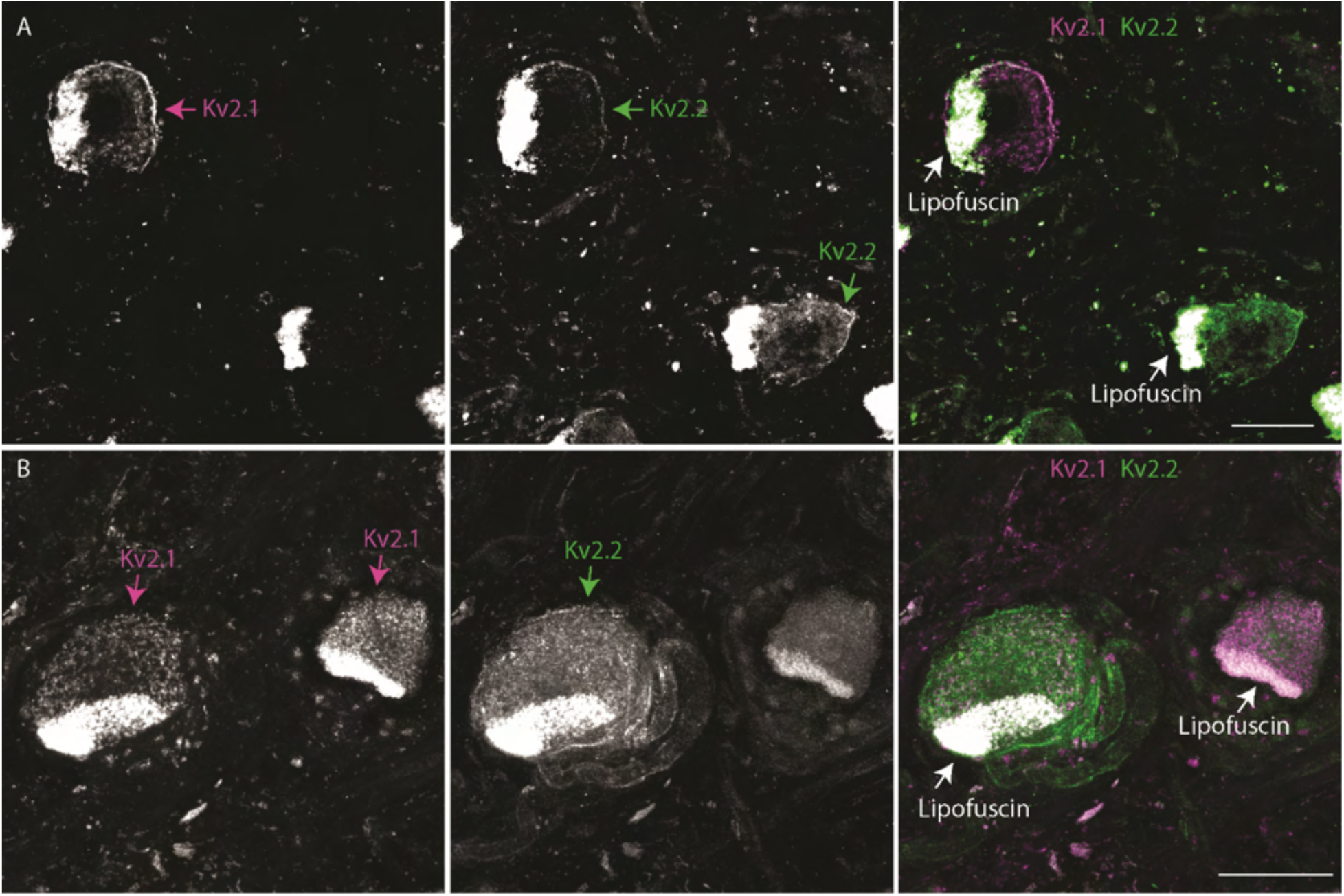
Immunofluorescence from human DRG neurons from donor #2 ***A*** and donor #3 ***B*** labeled with anti-Kv2.1 and anti-Kv2.2 antibodies. Autofluorescence attributed to lipofuscin is labeled in right panels while apparent Kv2.1 and Kv2.2 protein are labeled in left and middle panels respectively. Scale bars are 50 μm. Detailed information on each donor can be found in the *Human Tissue Collection* section of the methods.

**Supplemental Figure 13.**
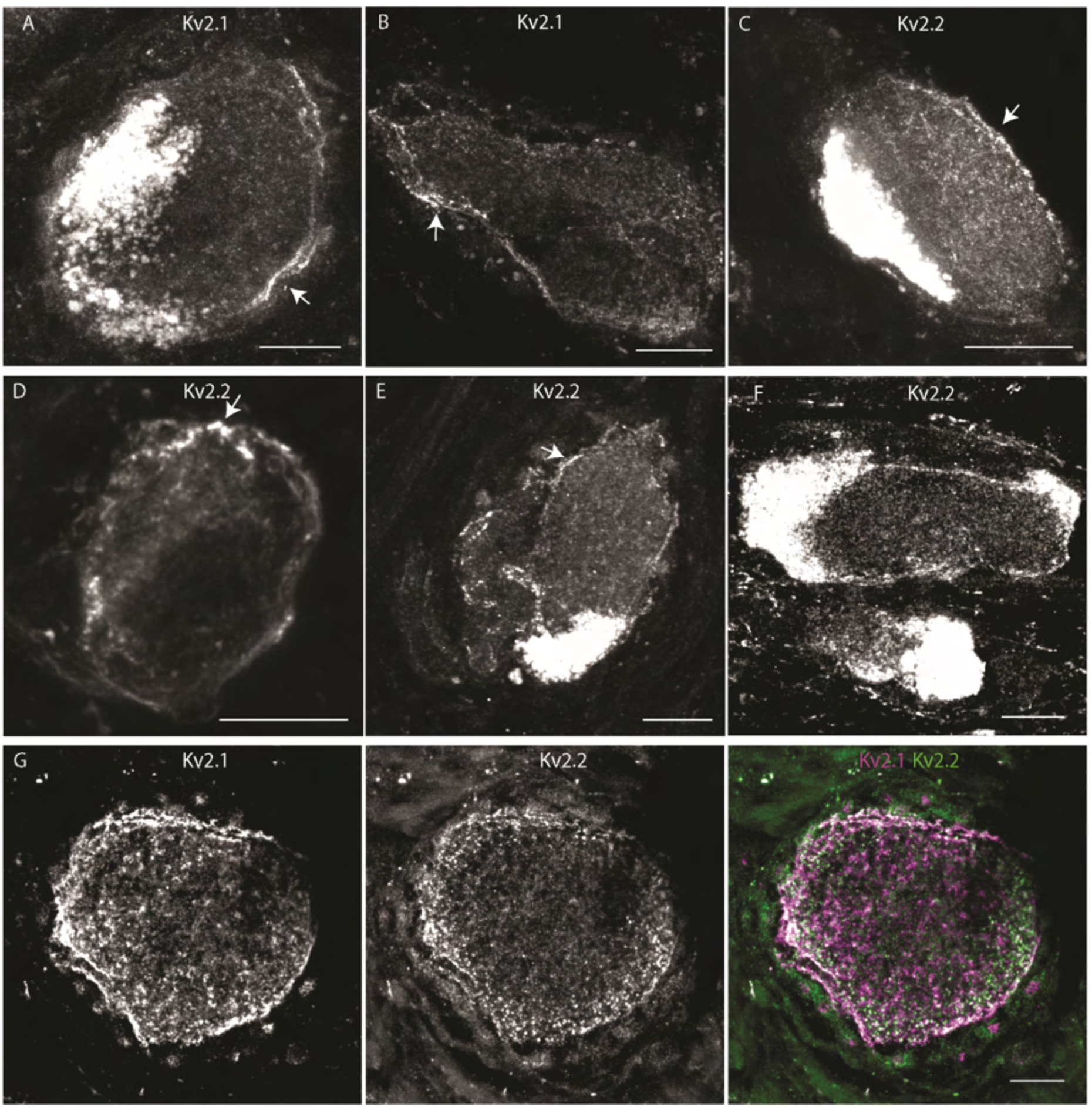
Kv2 channels are enriched at the outer edge of human DRG neurons. ***A-B***, Exemplar z-projections of anti-Kv2.1 immunofluorescence enriched at the outer surface of human DRG neurons. Arrows indicate asymmetric clusters. Images are from donor #2 Scale bars are 20 μm. ***C-F***, Exemplar z-projections of anti-Kv2.2 immunofluorescence enriched at the outer surface of a human DRG neurons. Arrows indicate asymmetric clusters. Image in ***E*** is from donor #3 while all other images are from donor #2. Scale bars are 20 μm. ***G***, Exemplar z-projection of anti-Kv2.1 and anti-Kv2.2 immunofluorescence both enriched at the outer surface of a human DRG neuron soma. Image is from donor #2. Scale bar is 20 μm. Detailed information on each donor can be found in the *Human Tissue Collection* section of the methods.

**Supplemental Figure 14.**
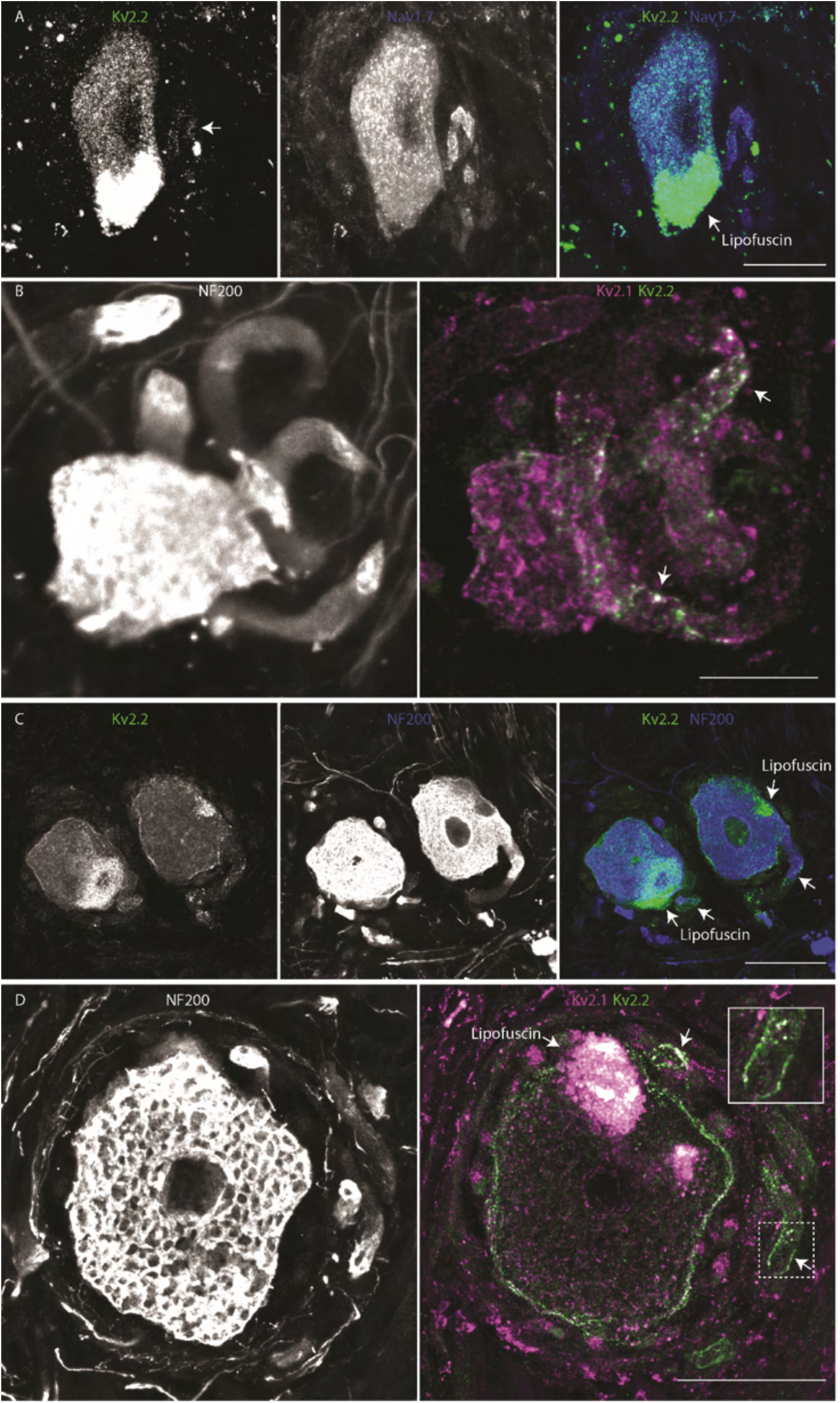
***A***, Z-projection of anti-Kv2.2 and anti-Nav1.7 immunofluorescence in a human DRG neuron soma and stem axon. Arrow in merge indicates the stem axon of the DRG neuron. Apparent lipofuscin autofluorescence is labeled in merge. Image is from donor #2. Scale bar is 20 μm. ***B,*** Z-projection of anti-Kv2.1 (magenta), anti-Kv2.2 (green) (right) and anti-NF200 (left) immunofluorescence in a human DRG neuron soma and stem axon. Arrows in merge indicate the stem axon of the DRG neuron. Image is from donor #1. Scale bar is 20 μm. ***C,*** Z-projection of anti-Kv2.2 (left) and anti-NF200 (middle) immunofluorescence in a human DRG neuron soma and stem axon. Arrow in merge indicates the stem axon of the DRG neuron. Apparent lipofuscin autofluorescence is labeled in merge. Image is from donor #1. Scale bar is 50 μm. ***D,*** Z-projection of anti-Kv2.1 (magenta), anti-Kv2.2 (green) (right) and anti-NF200 (left) immunofluorescence in a human DRG neuron soma and stem axon. Arrows in merge indicate the stem axon of the DRG neuron. Apparent lipofuscin autofluorescence is labeled in merge. Image is from donor #3. Scale bar is 50 μm. Detailed information on each donor can be found in the *Human Tissue Collection* section of the methods.

**Supplemental Figure 15.**
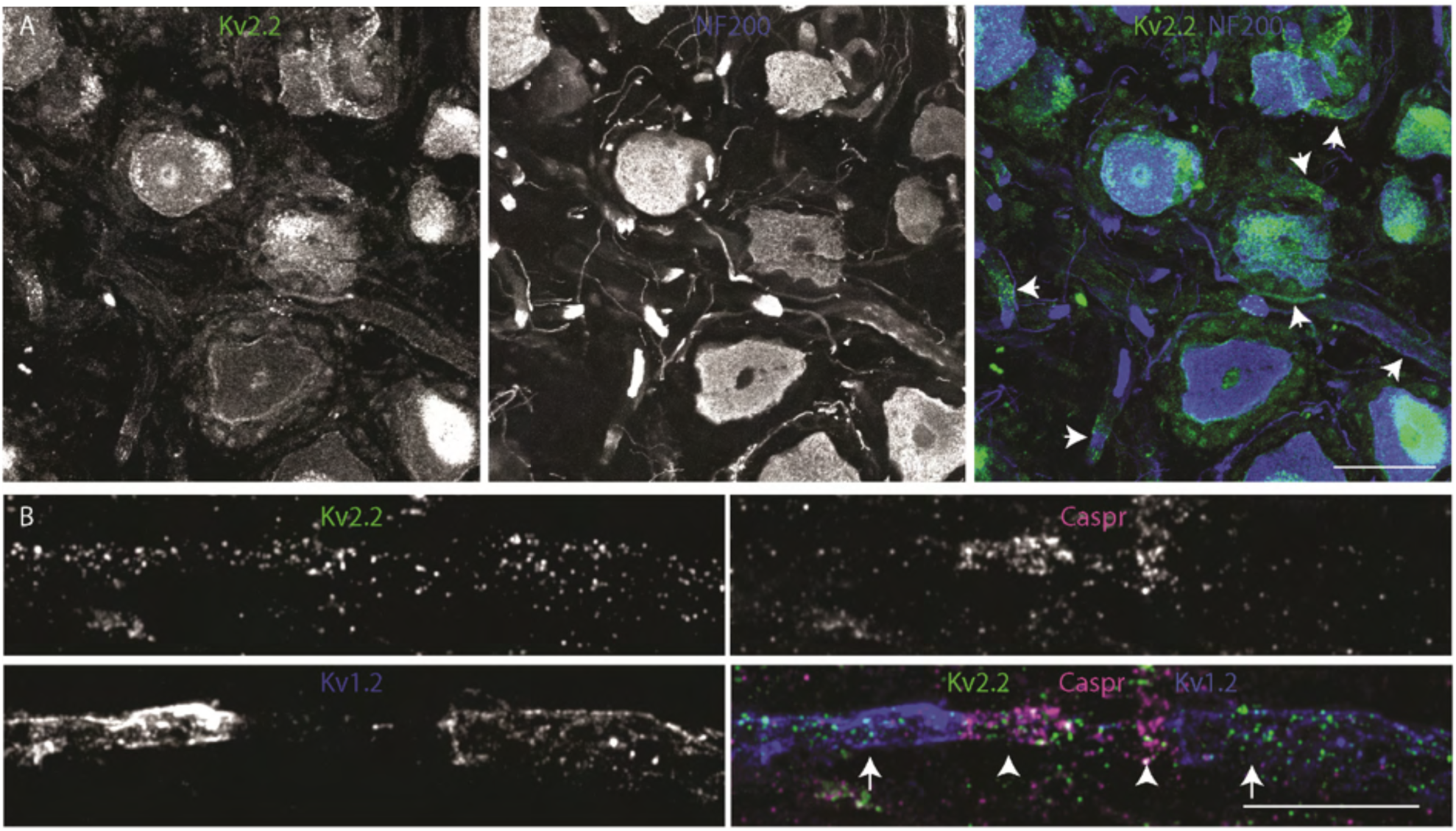
***A***, Z-projection of anti-Kv2.2 (left) and anti-NF200 (middle) immunofluorescence of human DRG. Arrows in merge represent exemplar axons which have clear anti-Kv2.2 immunofluorescence. Image is from donor #1. Scale bar is 50 μm. ***B***, Z-projection of anti-Kv2.2 (upper left), anti-CASPR (upper right) and anti-Kv1.2 (bottom left) immunofluorescence of human DRG axon. Image is from donor #2. Scale bar is 10 μm. Detailed information on each donor can be found in the *Human Tissue Collection* section of the methods.

